# Host-Pathogen Interactions in the *Plasmodium*-Infected Mouse Liver at Spatial and Single-Cell Resolution

**DOI:** 10.1101/2023.12.22.573046

**Authors:** Franziska Hildebrandt, Miren Urrutia-Iturritza, Christian Zwicker, Bavo Vanneste, Noémi Van Hul, Elisa Semle, Tales Pascini, Sami Saarenpää, Mengxiao He, Emma R. Andersson, Charlotte L. Scott, Joel Vega-Rodriguez, Joakim Lundeberg, Johan Ankarklev

**Author notes:** Corresponding authors: Franziska Hildebrandt & Johan Ankarklev, Department of Molecular Biosciences, The Wenner Gren Institute, Stockholm University, Svante Arrhenius väg 20C, SE-106 91, Stockholm, Sweden.

## Abstract

Upon infecting its vertebrate host, the malaria parasite initially invades the liver where it undergoes massive replication, whilst remaining clinically silent. The spatial coordination of factors regulating immune responses and metabolic zonation during malaria infection, in the true tissue context, remains unexplored. Here, we perform spatial transcriptomics in combination with single-nuclei RNA-sequencing (snRNA-seq) over multiple time points during liver infection to delineate transcriptional programs of host-pathogen interactions across *P. berghei-*infected liver tissues. Our data suggest changes in gene expression related to lipid metabolism in response to *Plasmodium* infection in the proximity of infected hepatocytes, such as the modulation of the expression of genes involved in peroxisome proliferator-activated receptor pathway signaling. The data further indicate the presence of inflammatory hotspots with distinct cell type compositions and differential liver inflammation programs along the lobular axis in the malaria-infected tissues. Furthermore, a significant upregulation of genes involved in inflammation is observed in liver tissues of control mice injected with mosquito salivary gland components, which is considerably delayed compared to *P. berghei* infected mice. Our study establishes a benchmark for investigating transcriptome changes during host-parasite interactions in tissues, it provides informative insights regarding *in vivo* study design linked to infection, and provides a useful tool for the discovery and validation of *de novo* intervention strategies aimed at malaria liver stage infection.

## INTRODUCTION

Infectious *Plasmodium spp.* sporozoites, transmitted by female *Anopheles* mosquitoes, escape the dermis after a mosquito bite and disseminates through the circulation, eventually infecting a liver hepatocyte ^1^. Inside the hepatocyte, the parasite forms a parasitophorous vacuole to obtain nutrients for growth and merozoite production ^2,3^. The parasite transitions into the symptomatic blood-stage by releasing thousands of merozoites ^4^ from an infected hepatocyte at around 48 hours post infection (hpi) for the rodent specific *Plasmodium berghei* parasite ^5^. Notably, the liver represents a major bottleneck during the malaria life cycle and is the stage targeted by the only WHO-recommended malaria vaccine to date. Despite the limited efficacy (36% in children 5-17 months of age ^6^) of the RTS,S vaccine, the pre-erythrocytic stages of malaria infection show substantial promise for further vaccine development.

The liver serves as a critical immune organ, detecting and eliminating pathogens and toxins while simultaneously regulating energy, lipids, and protein synthesis ^7,8^. Its structural organization consists of lobules, including hexagonal units with portal veins at the corners and a central vein at the center, making up metabolic zones, which is commonly referred to as zonation ^9,10^. Labor is further divided amongst the highly diverse cell types of the liver, including parenchymal cells, such as hepatocytes and cholangiocytes which account for 70 - 80% of the total liver area, as well as non-parenchymal cells (NPCs). NPCs include liver sinusoidal endothelial cells (LSECs), which line the vasculature of the liver, as well as Kupffer cells and other immune cells, including neutrophils, mononuclear cells, T and B lymphocytes, natural killer (NK) cells and NKT cells, which are found scattered across hepatic lobules ^11,12^. The portal vein is considered the main entry point of gut-derived pathogens making the liver susceptible to circulating pathogens ^7,12^. Maintaining immune balance is crucial for liver function, as disturbed homeostasis or prolonged inflammation can lead to severe diseases like cirrhosis, non-alcoholic steatohepatitis, hepatocellular carcinoma, and liver failure ^13^. However, pathogens like *Plasmodium* may exploit the liver’s immune tolerance ^14^.

During liver infection, *P. berghei* elicits a sequential transcriptional response, including interferon-mediated immune genes expressed at later parasite developmental stages in the liver ^15–17^. Parasite development in the liver is heterogeneous and suggested to be affected by zonation, where abortive infections in periportal zones have been described ^18^. These findings have advanced our understanding of *Plasmodium* infection and hepatocyte zonation, as well as tissue-wide immune responses. However, a comprehensive map of spatial host-parasite interactions, including gene expression profiles in their true tissue context, beyond hepatocyte zonation, and including the involvement of liver resident immune cells, has been missing.

In our previous work, we established the first spatial transcriptomics map of murine liver tissue, including expression by distance measurements of target structures ^19^. Here, we perform spatial gene expression analysis of *P. berghei-*infected mouse livers over multiple time points during infection (12-, 24- and 38-hours post-infection (hpi)) to map out genes and genetic pathways involved in host-parasite interactions across liver tissue sections. In this study we use a combination of the original Spatial Transcriptomics 2K arrays ^19,20^ (henceforth referred to as ST) and Visium (10X Genomics Inc.) ^21^. Spatial data resulting from ST enabled us to investigate a large sample size (n=38 tissue sections), whereas the Visium arrays (n = 8 tissue sections) allowed for increased resolution of expression analysis due to the decreased spot-size (55 µm *vs.* 100 µm) and shorter spot-center to center distances ^21^. Additionally, we performed single-nuclei RNA sequencing (snRNA-seq) on the same tissue samples to identify and deconvolve cell types. This integrated approach allows for a comprehensive transcriptomics analysis of *P. berghei*-infected liver sections, including complete cell type information.

Combining spatial transcriptomic and snRNA-seq data reveals both global and local effects of *P. berghei* infected liver tissue compared to controls. Notably, we identify differential expression of genes involved in lipid homeostasis at infection sites, potentially indicating a parasite immune evasion strategy. We also uncover unique tissue structures termed inflammatory hotspots (IHSs) that exhibit morphological and transcriptional distinctions and resemble focal immune cell infiltrates observed in liver pathologies of various diseases ^22–24^. Based on gene expression and cell type profiles, we propose that IHSs are sites of mechanical damage due to parasite traversal or sites of successful parasite elimination. In total, this study provides a highly informative resource of spatio-temporal host tissue responses during malaria infection and development in the liver.

## RESULTS

### Spatial Transcriptomics captures liver tissue responses induced by malaria parasite infection

We used Spatial Transcriptomics (ST)^19^ to analyze 38 liver sections of 18 adult female mice, infected with either *P. berghei* parasites or uninfected *An. gambiae* salivary gland lysate (SGC) at different time points (12, 24, and 38 hours post-infection). We added Visium Spatial Gene Expression experiments for higher spatial resolution (see methods for details), resulting in a total of 46 spatially analyzed liver sections. The SGC sections helped control for mosquito-related responses. In addition, we performed single-nuclei RNA-sequencing (snRNA-seq) to deconvolve spatial data and increase the resolution in our analyses further (Figure 1a).

**Figure 1.**
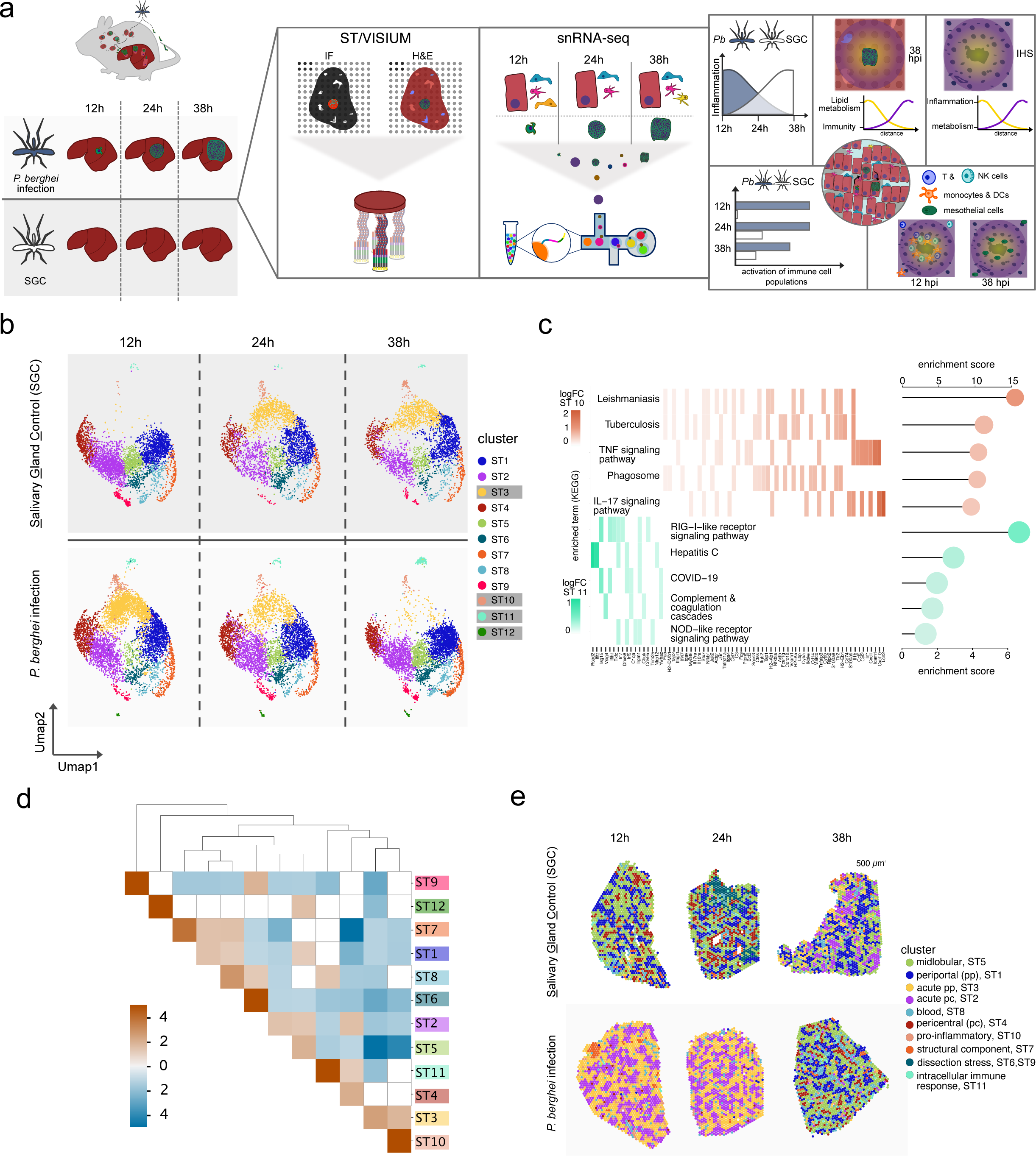
Spatial organization of livers infected with *P. berghei* parasites or SGC. **a)** Schematic representation of experimental design of this study. Livers were collected at 12, 24 or 38 hpi with *P. berghei* parasites or salivary gland lysate of uninfected mosquitoes (SGC) (left). Immunofluorescence staining of the parasite and ST or 10X Visium spatial technology protocols were performed. Simultaneously, droplet-based single nuclei RNA sequencing (snRNA-seq) was performed for all infection conditions (center). Both data were further analyzed computationally, for example including spatial as well as cell clustering and annotation based on expression profiles, expression by distance analysis and differential gene expression analysis (DGEA) between infected (INF.) and SGC (C). **b)** Liver sections from ST analysis were normalized and batch-corrected. After dimensionality reduction the data was embedded in UMAP space and split by the original condition for visualization. Data from SGC sections are shown on the top from 12-38 hpi (left to right) and data from *P. berghei* infected sections are shown on the bottom from 12-38 hpi (left to right). Clusters with an obvious association to infection condition are highlighted with gray boxes in the legend. **c)** For identified clusters ST10 and ST11, differential genes expression analysis (DGEA) was performed followed by functional enrichment analysis for each cluster (see methods for details). Overrepresented pathways of the KEGG database for ST10 are shown in rose and for ST11 in aquamarin. Scales for expression values for overrepresented genes belonging to the individual KEGG pathways are shown for ST11 (left) or ST10 (right), from high expression (dark) to lower expression (light). Selected gene names are shown at the bottom. Enrichment scores for the pathways are shown on the right. **d)** Interaction analysis of clusters was performed to evaluate spatial enrichment expression programs as suggested by clustering analysis in space. Positive enrichment values (orange) indicate spots belonging to these clusters are more likely to be neighboring while negative enrichment values (blue) indicate spots associated with these expression programs are less likely to be neighboring. Clusters without significant enrichment in each other’s neighborhoods are shown in white. **e)** 10X Visium experiments were performed in the same fashion as ST experiments and clustering generated similar results. Clusters were imposed on spatial positions and annotated according to spatial expression features. Sections of the investigated conditions are divided for ease of inspection as in **b)**, with the top panel comprising SGC sections across 12 -38 hpi (top, left to right) and the bottom panel comprising *P. berghei* infected sections across 12 - 38 hpi (bottom, left to right).

We first identified spatial expression patterns related to infection by performing unsupervised clustering analysis (see methods for details). We identified 12 clusters for the ST data (ST1 - ST12) (Figure 1b, Supplementary figures 1-3) and 10 clusters for the 10X Visium data (Figure 1c). Four of these ST clusters - namely ST3, ST10, ST11, and ST12 - exhibited a unique pattern of gene expression influenced by the condition i.e. *P. berghei* infection or SGC challenge, and the collection time point (12h, 24h, or 38h) (Figure 1b, Supplementary figure 4). At 12 hpi with *P. berghei*, a large proportion of spots displayed ST3 expression, while SGC-challenged mice showed the opposite trend, but with increasing proportions of ST3 at later time points. There was a similar observation for ST10, but with fewer associated spots. Spots belonging to ST11 show enrichment in sections infected with *P. berghei* parasites while spots of ST12 are missing entirely from SGC sections (Figure 1b, Supplementary figure 4).

Differential gene expression analysis (DGEA) revealed that cluster ST12 is defined by upregulation of *P. berghei* specific transcripts (*HSP70-pb*, *HSP90-pb*, *LISP2-pb*), suggesting they represent parasite infected tissue sites (Supplementary figure 5, Supplementary data 1). Spots associated with clusters ST10 and ST11 exhibit an anti-correlated presence along the infection timeline. Further, DGEA and gene ontology (GO) enrichment suggests that ST10, predominantly active during early infection, is associated with pro-inflammatory signaling (e.g., IL-17 and TNF pathways), including phagocytosis, and KEGG-terms including leishmaniasis and tuberculosis. In contrast, ST11 is enriched in pathways related to intracellular pathogen signaling (NOD-like and RIG-I-like receptor pathways), complement and coagulation cascades, and KEGG terms associated with viral infections such as COVID-19 and Hepatitis C. Moreover, most upregulated genes in ST11 are interferon-stimulated genes (ISGs), including *Ifit1*, *Ifih1*, *Irf7*, and *Irf9* (Figure 1d, Supplementary figure 5, Supplementary data 1).

Differentially expressed genes (DEGs) in the remaining clusters include ST3, which exhibits an upregulation of genes linked to acute phase response and inflammation, including the *Saa* ^25,26^ and *Orm* families (Supplementary figure 5, Supplementary data 1). The higher prevalence of ST3 associated spots at 12 hpi suggests an initial inflammatory stress response in the *P. berghei* infected liver, which is delayed in the SGC sections.

Several of the identified clusters (ST1, ST4-ST5, and ST7-ST8) were previously described in healthy liver tissue and validated here ^19^. These clusters represent periportal, pericentral, midlobular zonation, structural integrity, and blood cell populations (Figure 1b).

We identified three new clusters (ST2, ST6, and ST9) with previously undescribed expression profiles. These clusters do not show clear links to *P. berghei* infection or SGC challenge (Figure 1b, Supplementary table 1). Cluster ST2 exhibits expression of a number of genes which are associated with pericentral localization, such as *Cyp2e1* ^10,19,27^ (Supplementary figure 5), suggesting it may represent an intermediate zone between central and portal areas, closer to the central region. We confirmed this by analyzing cluster interactions, showing that cluster ST2 is enriched in spots adjacent to cluster ST4 (Figure 1e), supporting its pericentral proximity.

Comparing ST and Visium data reveals significant overlap in differentially expressed genes (DEGs) across identified clusters (Supplementary figure 6, Supplementary data 1). Notably, spots associated with *P. berghei* infection (ST12) are not present in every analyzed infected tissue section. This is especially the case at the early infection time points where the number of detected *P. berghei* infected hepatocytes per tissue section is lower, emphasizing the value of larger sample sizes for ST experiments. Further, the higher resolution of Visium compared with ST enables the distinction of spatial gene expression patterns in clusters ST2 (acute pericentral) and ST3 (acute periportal) (Figure 1c), suggesting zonation of the acute response during infection.

### *P. berghei* infection impacts both proximal and peripheral gene expression in liver tissue

We found the majority of uniquely DEGs between *P. berghei* infected and SGC sections at 12 and at 38 hpi (Supplementary figure 7, Supplementary data 2). Upregulated genes at 12 hpi in *P. berghei* infected tissues are linked to cellular stress responses including transcription of *Saa1*, *Saa2*, *Saa3* and *Lcn2* ^26,28^. Meanwhile, most upregulated genes at 38 hpi belong to the group of ISGs including *Ifit1*, *Ifit3*, *Irf7* and *Usp18*, which have been previously implicated with an interferon response towards *Plasmodium* liver infection ^15,18,29^ (Figure 2a, Supplementary data 2).

**Figure 2.**
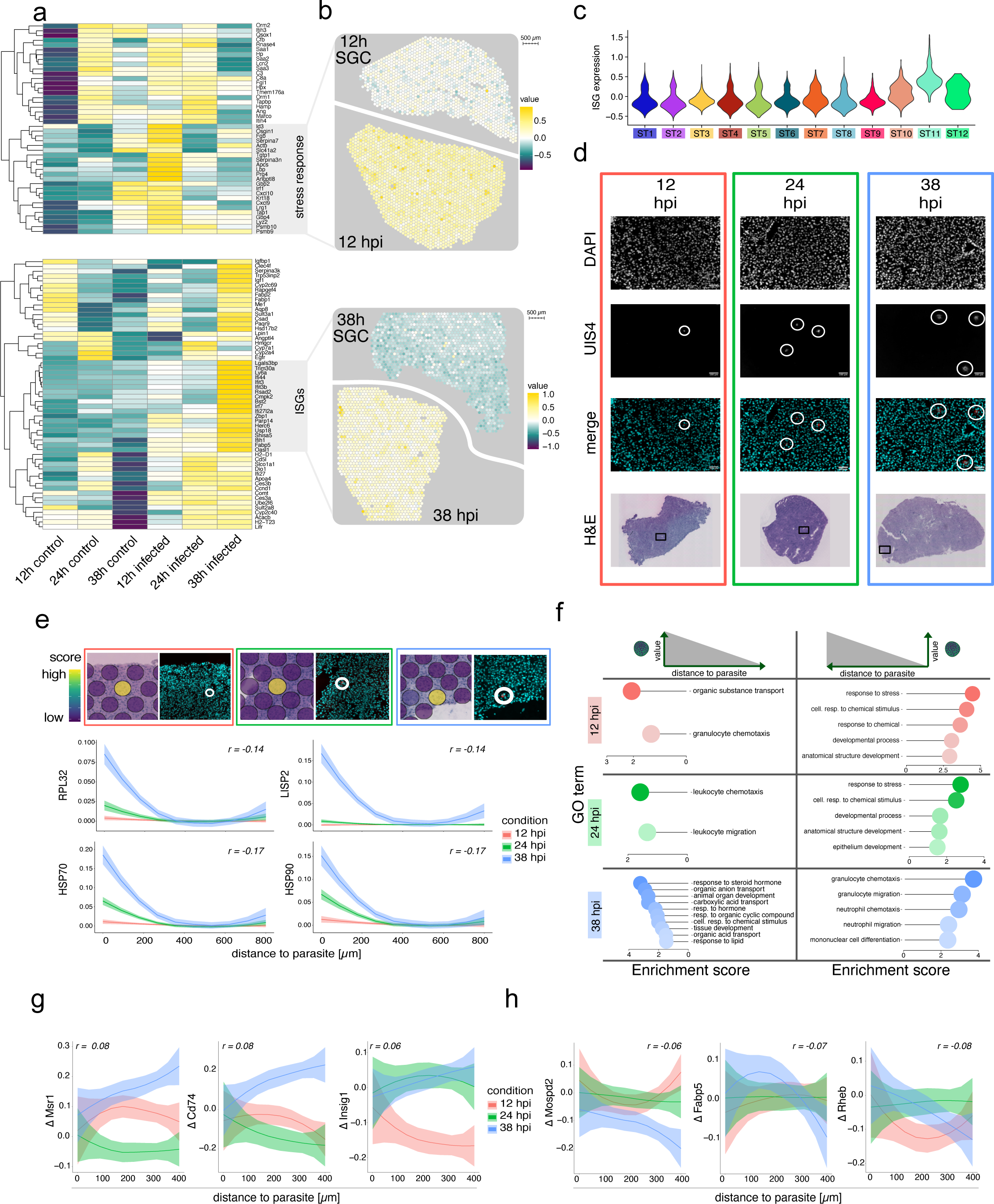
Global and spatially distinct effects of *P. berghei* on tissue gene expression. **a)** Heatmap of differentially expressed genes between *P. berghei* infected sections and SGC sections at 12 hpi (top) and 38 hpi (bottom). Genes are clustered hierarchically based on expression similarity. Averaged gene expression is shown as a color gradient from low (dark purple) to high (yellow) expression. Function of expression programs with highest upregulation in the respective time points are highlighted in gray boxes, comprising “stress response” at 12 hpi or “Interferon-stimulated genes (ISGs)” at 38 hpi. **b)** Expression of modules showing highest expression values in 12 hpi (stress response) and 38 hpi (ISGs) across tissue spots for infected and SGC tissue sections at the respective time points. The scale bar denotes 500µm and module expression values are depicted as a scale ranging from low expression (dark purple) to high expression (yellow). **c)** Violin plot showing expression of ISGs module across spatial clusters. Clusters are depicted in the same colors as previously established in Figure 1a. **d)** Immunofluorescence and Hematoxylin and Eosin (H&E) stained images of *P. berghei* infected tissue sections across investigated conditions at 12h, 24h and 38h (left to right). Colored boxes indicate time points (12 hpi = red, 24 hpi = green, 38 hpi = blue). Positions with parasites are shown from individual IF images, showing DNA staining (DAPI), parasite staining (UIS4) and the composite image (merge). Parasites are highlighted by white circles and scale bars indicate 100µm. The position of detected parasites is shown as a black box on the respective H&E images, recorded after immunofluorescent staining. **e)** Visualization of module scores of top *P. berghei* genes with negative correlation to parasite distance (top). Colored boxes indicate time points (12 hpi - 38 hpi, left to right) as in **d)** and white circles indicate positive parasite signal (UIS4). Module scores on corresponding H&E images show high expression as a scale from low (dark purple) to high (yellow). Expression-by-distance analysis of *P. berghei* genes with negative correlation to parasite distance shows change of expression values as a function of the distance between 0 and 800 µm from parasite neighborhoods (methods for details) at 12, 24 and 38 hpi. Correlation values are indicated by r. **f)** Gene-Ontology (GO) enrichment of top five GO-terms of genes associated with close distance to the parasites (left) or far distance to the parasite (right). Colors indicate time points as in **d)** (12 hpi -38h hpi, top to bottom). **g)** Change in gene expression (Δ) of selection of host genes exhibiting negative correlation to distance to parasite neighborhoods (associated with close proximity to parasite) within 400 µm to parasite neighborhoods across time points of infection. **h)** Change in gene expression (Δ) of selection of host genes exhibiting positive correlation to distance to parasite neighborhoods (associated with further proximity to parasite) within 400 µm to parasite neighborhoods across time points of infection.

Modules of stress response genes at 12 hpi and ISGs at 38 hpi exhibited higher expression in infected sections, but this expression was not confined to the infection sites, suggesting a widespread inflammatory response across the tissue (Figure 2b). Cluster ST11 displayed the highest expression of ISGs, indicating that the location of cluster ST11 represent foci of type I IFN response (Figure 2c).

Unsupervised clustering results indicate parasite localization across the infected tissues. However, determining parasite positions solely at the RNA level proves challenging due to limited spatial resolution and low parasite transcript abundance. Despite these challenges, we can detect an increased number of parasite transcripts in the infected conditions over time (Supplementary figure 8). In addition, robust validation of parasite positions and development is achieved through immunofluorescence (IF) staining using the parasitophorous vacuole membrane (PVM) marker UIS4 (Figure 2d).

We performed a correlation analysis between the distance to the neighborhood of the parasite annotation and gene expression (see methods for details). To facilitate the interpretation of expression changes (Δ) across conditions, we centered expression at 0 µm from the parasite neighborhood. Negative correlation signifies reduced expression with increased proximity to the parasite, while positive values indicate increased expression. Notably, we observed a significant negative correlation between parasite distance and overall parasite gene expression within 400 µm of parasite neighborhoods, peaking at 38 hpi (Figure 2e).

Despite the significantly lower abundance of parasite transcripts compared to the host, we performed DGEA in parasite neighborhoods, aligning it with Afriat et al.’s pseudotime analysis ^18^. This revealed a high proportion of genes from our data is linked to early latent time determined by Afriat et al., which is possibly due to the sparse presence of *P. berghei* transcripts in our data (Supplementary Figure 9).

Next, we determined host gene expression with positive and negative correlation to *P. berghei* infection sites across all time points and performed a GO term enrichment analysis (see methods for details). The GO term enrichment suggests higher expression of genes involved in chemotaxis of leukocytes, including expression of *Xcl1*, *Fcer1g* and *Csf1r* near the parasite at 12 and 24 hpi. However, the pattern is reversed at 38 hpi, with decreased expression of the genes described above, along with other genes including, *Msr1*, *Cd74*, *Csf3r*, and *Camk1d*. These genes exhibit higher expression with increased distance from the infection site (Figure 2f-g, Supplementary figure 10-13, Supplementary data 3).

Leukocyte chemotaxis is crucial for inflammation and immune responses and includes the recruitment of macrophages and neutrophils to ward off invading pathogens ^30,31^. Our data indicate that the parasite triggers a pro-inflammatory response near the infection site but evades phagocytosis during the late infection time-point, just prior to egress from the liver. Notably, at 38 hpi, genes such as *Msr1* and *Cd74*, which are associated with inflammation ^32,33^, show positive correlation with increasing distance from the parasite (Figure 2g, Supplementary Figure 12, Supplementary Data 3). Additionally, we found *Insig1*, which is linked to lipid homeostasis and the prevention of lipid toxicity ^34^, to positively correlate with parasite neighborhoods (Figure 2g, Supplementary Figure 12, Supplementary Data 3).

In the proximity of parasite locations, we observed higher expression of *Fabp5*, involved in the regulation of lipid metabolism, peroxisome proliferator-activated receptors (PPARs) and cell growth ^35,36^. We also identified higher expression of *Mospd2*, implicated in host-pathogen interactions with *T. gondii* ^37^, and *Rheb* expression close to parasites (Figure 2h, Supplementary Figure 13, Supplementary Data 3). Rheb activates mTORC1, promoting proliferation and survival. Moreover, Rheb is shown to inhibit autophagy ^38–40^, an increasingly recognized pathway in *Plasmodium* liver infection ^41,42^.

### Inflammation exhibits spatial patterns in response to *P. berghei* and SGC challenge

Several studies implicate that parasite localization in the different metabolic zones of the liver influences the developmental progress of *Plasmodium* in hepatocytes and suggest higher developmental success in pericentral areas ^18,43,44^. While our data do not indicate direct correlation between hepatic zonation and *P. berghei* localization in liver tissue, we observe similar trends, where parasite gene expression is higher in areas within 400 µm of computationally annotated pericentral veins (see methods for details). In addition, our data suggest that a large proportion of parasites are present and transcriptionally active in areas that we defined as intermediate, situated beyond 400 µm from both pericentral and periportal neighborhoods (Supplementary figure 14).

Our data further suggest hepatic zonation of inflammatory responses at 12 and 24 hpi (Figure 1e). To further validate this observation, we investigated correlations between periportal marker genes (*Cyp2f2*, *Sds*), pericentral marker genes (*Glul*, *Slc1a2*) and differentially expressed genes in the acute periportal cluster (ST3) or the acute pericentral cluster (ST2). Marker genes of ST2 (*Car3*, *Ces3a*, *Ces1d*, *Cyp3a11*, *Nr1i3*) correlate with gene expression of pericentral marker genes while marker genes of ST3 (*Itih3*, *Itih4*, *C3*, *Ambp*, *Fgg*, *Qsox1*, *Hpx*) correlated with periportal marker genes (Figure 3a), supporting the notion of hepatic zonation. Expression-by-distance analysis further validated zonated expression profiles of acute periportal and acute pericentral genes (Figure 3b).

**Figure 3.**
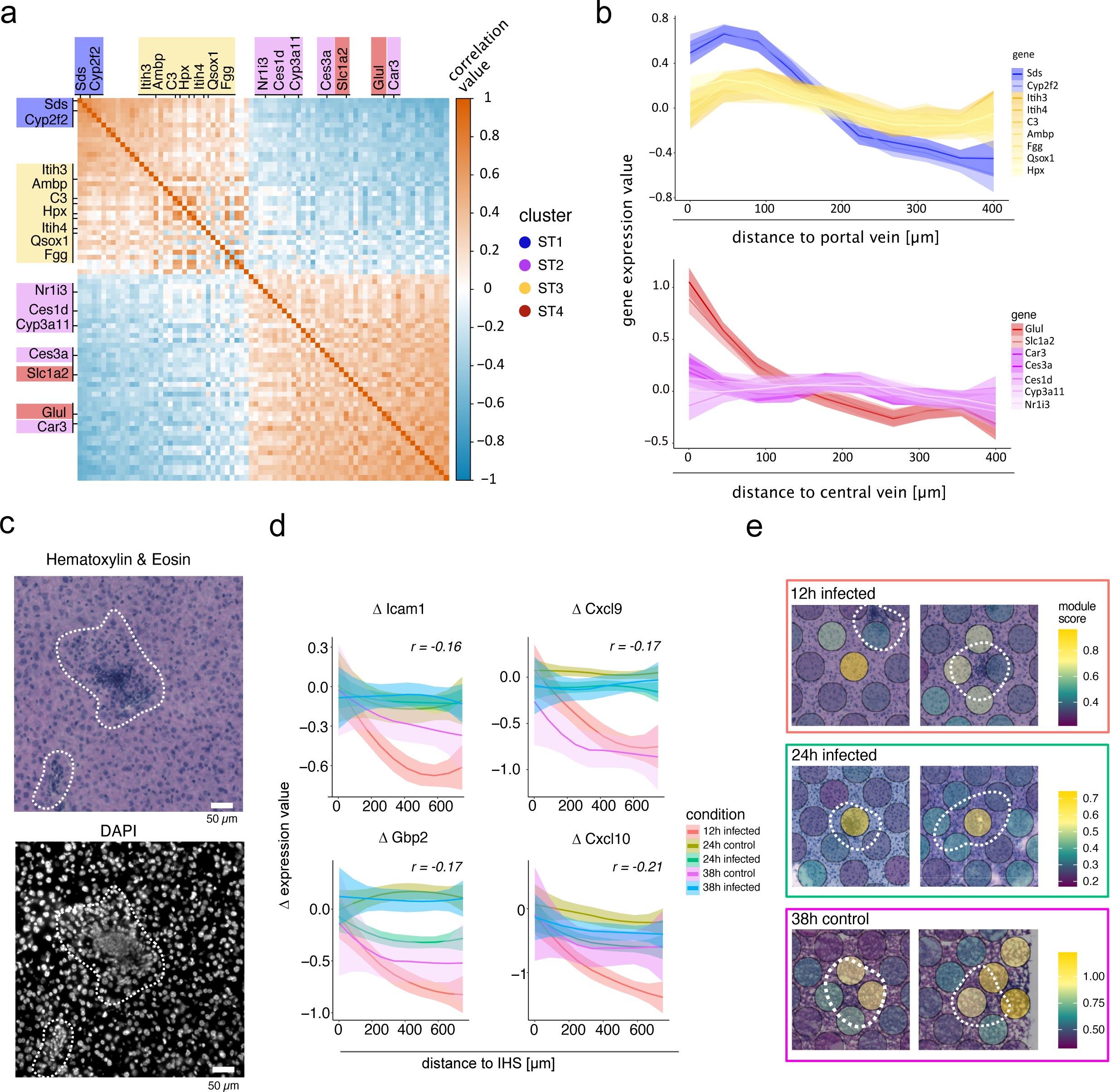
Spatial inflammation in *P. berghei* infected and SGC sections. **a)** Pearson correlations between marker genes of spots belonging to periportal in cluster ST1 (blue), acute inflammation in cluster ST3 (yellow), pericentral in cluster ST4 (red) and acute pericentral in cluster ST2 (purple). Positive correlation values are indicated in orange and negative correlation values are indicated in blue. **b)** Gene expression of genes highlighted in **a)** as a function of the distance to the portal vein for marker genes of cluster ST1 and ST3 (top) or the central vein for marker genes of cluster ST2 and ST4. **c)** Representative H&E (top) and DAPI (bottom) images of Inflammation hotspots (IHSs) observed in *P.berghei* infected section 12 hpi. IHSs are highlighted with white dotted lines. The scale bar indicates 50 µm. **d)** Change in expression (Δ) of top 4 genes with highest negative correlation as a function of the distance between 0 and 600 µm from IHSs neighborhoods (methods for details) where IHSs were present (12, 24 and 38 hpi as well as 24 and 38 hours after salivary gland challenge (control)). **e)** Projection of expression modules of genes in **d)** on tissue sections across three conditions with highest numbers of visually annotated IHSs (12 and 24 hpi as well as 38h after salivary gland challenge (control)). Module scores are shown as color gradient from low scores (dark purple) to high scores (yellow). IHSs are highlighted with white dotted lines. View fields measure 500 by 500 µm.

Together with our observation that parasite numbers are increased in intermediate regions of hepatic zonation, this observation suggests that zonated inflammatory response to a high dose-infection may influence parasite survival and assist potential clearance, both in periportal and pericentral areas.

In addition to zonated inflammation, our data suggest a delayed global inflammatory response in SGC-challenged mice compared to *P. berghei* infection. Histological annotations reveal immune cell infiltration resembling focal structures, characterized by increased DNA signal (Figure 3c). These structures, which we have termed inflammatory hotspots (IHSs), follow the same trend as the global inflammatory response, primarily appearing at 12 and 24 hpi in the infected conditions and at lower frequency at 38 hpi. We explored gene expression profiles correlated with the distance from IHSs and found genes linked to inflammation and immune responses (Supplementary figure 15-16, Supplementary data 3). The four genes with the strongest negative correlation to IHSs include *Icam1*, *Gbp2*, *Cxcl9* and *Cxcl10* (Figure 3d-e).

Cxcl9 and Cxcl10 are key pro-inflammatory cytokines attracting activated T cells to inflammation sites ^45,46^. Gbp2 exhibits antiviral activity in murine macrophages and is upregulated during infection ^47^. Icam1 is upregulated by several cell types, including macrophages and regulates leukocyte recruitment from circulation to inflammation sites ^48^. Notably, Cxcl10 upregulation in infected hepatocytes is tied to the previously described abortive parasite phenotype ^18^. Additionally, IHSs seem to develop preferentially in periportal proximity (Supplementary figure 17).

### snRNA-seq and spatial integration reveal differential expression programs and suggest enrichment of various immune cell types in the IHSs

snRNA-seq enabled us to define distinct cell populations and their differential gene expression patterns across infection conditions and to further deconvolute cell type information of spatial gene expression data and estimate cell type proportions across the tissue.

Comparing proportions of 14 different annotated cell types (Figure 4a, Supplementary data 4), we find 70-80% hepatocytes and 20-30% remaining cell types (Supplementary figure 18). Cell type proportions of the 4 identified immune cell clusters (Kupffer cells, monocytes and DCs, T and NK cells and B cells) showed no significant difference in proportions between infected and SGC samples, at any time point but only trends of increased proportions of Kupffer cells, monocytes and DCs in infected conditions (Figure 4b).

**Figure 4.**
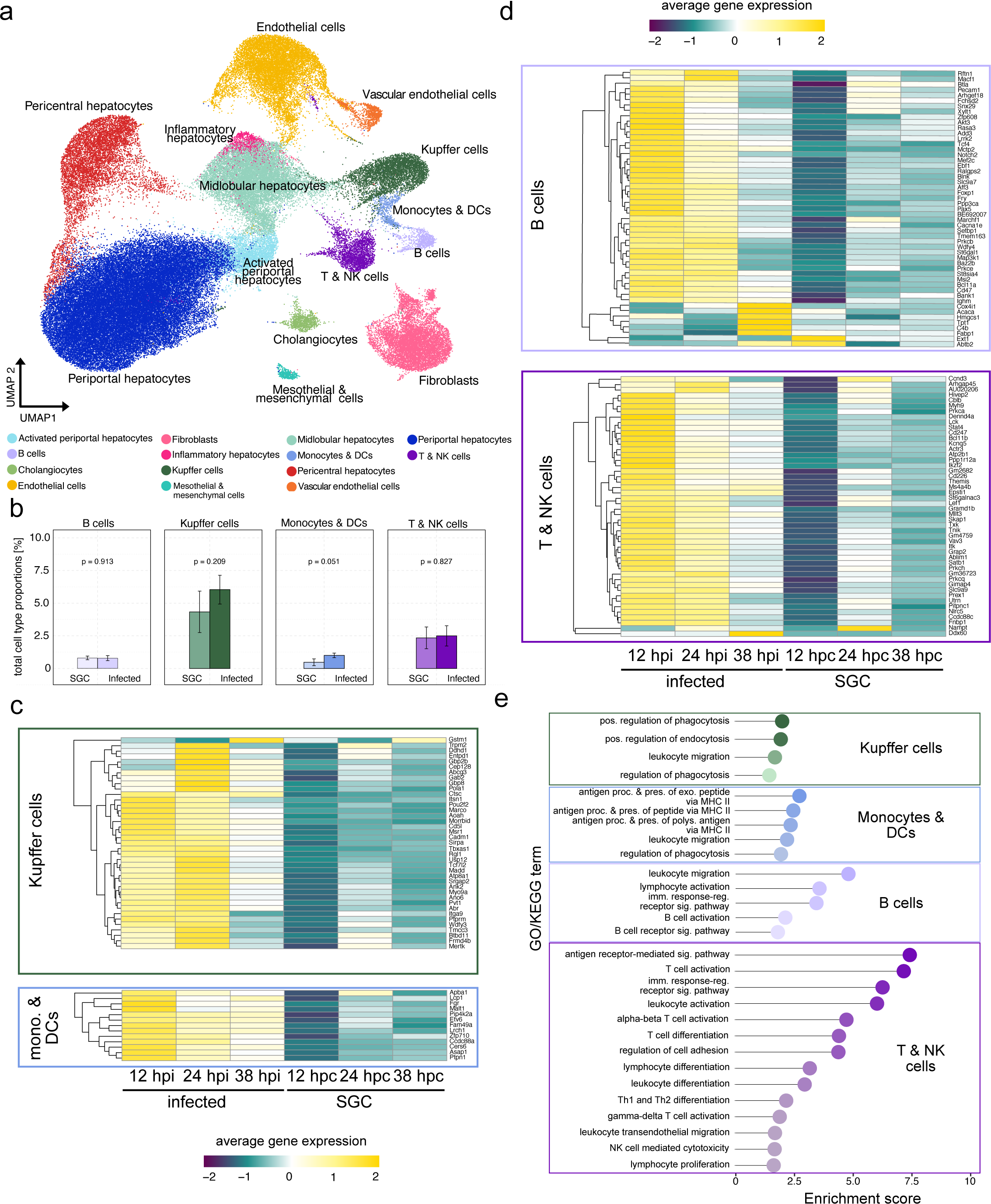
Identification of liver cell types and differential gene expression of immune cell clusters across infection conditions. **a)** UMAP projection of annotated liver cell types after integration of single cell expression data of all infection conditions: 12, 24 and 38 hpi as well as 12h, 24h and 38h post challenge with salivary gland lysate (SGC). **b)** Average immune cell type proportions normalized to the total number of different immune cells (T & NK cells, B cells, Monocytes & DCs and Kupffer cells) divided by the total number of cells. Error bars indicate the standard error of the mean across time points (see methods for details). **c)** Heatmap visualization of differential gene expression of genes associated with cell types of the myeloid lineage including Kupffer cells, Monocytes (mono.) and Dendritic cells (DCs) across infection conditions and time points. Average gene expression across respective cell types is depicted in a color scale ranging from high (yellow) to low (purple). **d)** Heatmap visualization of differential gene expression of genes associated with cell types of the lymphatic lineage including B cells, T cells and NK cells across infection conditions and time points. Average gene expression across respective cell types is depicted in a color scale ranging from high (yellow) to low (purple). **e)** Gene-Ontology (GO) enrichment of GO- or KEGG-terms of unique genes associated with different cell types. Colors indicate the respective immune cell type including Kupffer cells, Monocytes & DCs, B cells and T & NK cells (top to bottom).

We explored immune cell expression differences across conditions, noting upregulation of distinct genes for each immune cell type in infected livers at all time points compared to SGC controls (Figure 4c-d, Supplementary data 4). Infection-related marker genes within immune cell types exhibited higher expression at early time points (12 and 24 hpi), declining by 38 hpi. While expression in SGC controls increased over time, it did not reach the same levels as seen in infected cells (Figure 4c-d).

GO enrichment analysis revealed pathways associated with phagocytosis and leukocyte migration in Kupffer cells (e.g., *Marco*, *Msr1*, *Mertk*, *Cadm1*, *Itga9*, *Trpm2*). Monocytes and DCs were enriched for antigen presentation via MHC class II (*H2-Aa*, *H2-Eb1*, *H2-Ab1*, *Psap*). Lymphoid lineage cells (B, T/NK cells) showed enrichment in leukocyte migration (*Itk*, *Txk*), activation (*Bcl11a*, *Mef2c* for B cells; *Bcl11b*, *Satb1* for T cells), and NK-mediated cytotoxicity (*Cd247*, *Lck*, *Vav3*, *Prkca*) (Figure 4e). Thus, GO-term enrichment analysis, along with DGEA, confirms cell types and suggests their heightened activity in *P. berghei* liver infection.

The spatial organization of the different identified cell types across liver tissue sections confirmed the expected anti-correlated distribution of pericentral and periportal hepatocytes across tissue sections (Figure 5a, Supplementary figure 19). This was further validated by proportion-by-distance analysis (proportion-by-distance), using central or portal vein neighborhoods as the center (Figure 5b).

**Figure 5.**
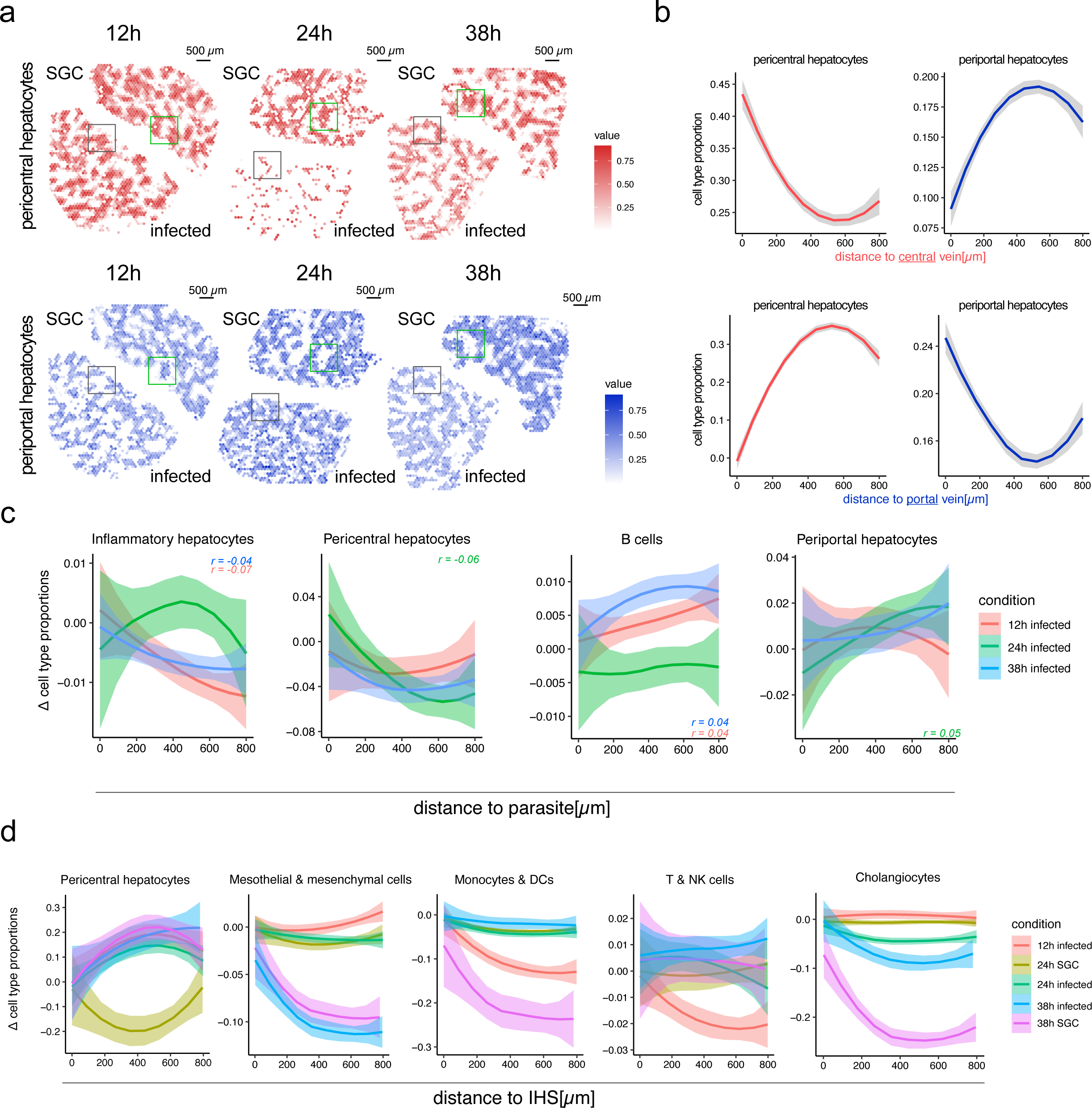
Integration of spatial and single nuclei data. **a)** Visualization of pericentral (top) and periportal (bottom) cell type proportions across spatial positions of sections generated by 10X Visium protocol. Pericentral cell type proportions are shown in red and periportal cell type proportions in blue. Green and gray boxes highlight smaller regions of opposite cell type compositions in salivary gland lysate control (SGC) and infected sections, respectively, for ease of inspection. The scale bars indicate 500 µm. **b)** Pericentral and periportal cell type proportions along a distance between 0 and 800 µm originating at computationally annotated central (top) or portal (bottom) veins. Periportal hepatocyte proportions are shown in blue and pericentral cell type proportions in red. Gray ribbons indicate standard error. **c)** Change in cell type proportions (Δ) of cell types with significant (p <= 0.05) negative (inflammatory hepatocytes, pericentral hepatocytes) or positive (B cells, periportal hepatocytes) correlation between the distance of 0 to 800 µm to parasite neighborhoods (methods for details). Conditions are indicated by colors (12h infected = red, 24h infected = green, 38h infected = blue). Correlation values (r) are indicated for each condition in the respective color. **d)** Change in cell type proportions (Δ) of cell types with significant (p <= 0.05) positive (pericentral hepatocytes) or negative (mesothelial & mesenchymal cells, T & NK cells, Monocytes & DCs) correlation between the distance of 0 and 800 µm to IHS neighborhoods (methods for details) where IHSs were present (12, 24 and 38 hpi as well as 24 and 38 hours after salivary gland challenge (control)). Correlations were calculated jointly for all time points.

Pearson correlations between cell type proportions and their distance to parasite neighborhoods across time points identified significant positive (indicating lower cell type proportions near the parasite) or negative (indicating higher cell type proportions near the parasite) correlations (Figure 5c). Cell types with increased proportions near the parasite included “inflammatory hepatocytes” at 12 and 38 hpi, characterized by stress response and inflammation markers (*Saa1, Saa2, Saa3, Ifitm3, Ly6e*) and a hepatocyte gene signature (*Alb, Apoc3, Apoh, Hamp, Cyp1e2*), as well as pericentral hepatocytes at 24 hpi. Conversely, cell types with decreased proportions near the parasite included B cells at 12 and 38 hpi and periportal hepatocytes at 24 hpi (Figure 5c).

Despite significant correlation, observed changes in cell type proportions relative to parasite neighborhood distance are small. This suggests that parasites may either have a minor impact on these cell type compositions in the liver tissue, or that only a few cells of these cell types are responsible for the observed differences.

Lastly, we established Pearson correlations between cell type proportions and distances to IHS neighborhoods, jointly analyzing all time points due to the limited number of IHSs. Positive correlations were observed for pericentral hepatocytes in all conditions except 24h SGC, while negative correlations were observed cholangiocytes at 38 hpi and in controls indicating a preference for IHSs to locate far from pericentral veins and closer to periportal areas (Figure 5d). Additionally, we noted higher proportions of T/NK cells and monocytes/DCs at IHSs in early infected (12hpi) sections and 38h SGC sections (Figure 5d). These cell types play critical roles in the immune response, as they produce various cytokines and communicate through cytolytic mechanisms ^49^. To characterize these cell infiltrates further, we employed IF staining, revealing increased lymphocytic (CD4, CD8) and myeloid cell (CD11b) infiltration and activation over time in infected and control livers, albeit delayed in SGC-treated mice (Supplementary Figures 20-22). F4/80+ macrophages within IHSs exhibited the highest abundance at 24h in infected livers and 38h in control livers (Supplementary figures 20-22). Notably, CD27 was exclusively detectable in *P. berghei*-infected livers at all time points, indicating heightened lymphocyte activation compared to controls (Supplementary Figures 20-22) ^50^. Together with previous studies, where higher proportions of extracellular matrix producing mesothelial and mesenchymal cells have been described ^51,52^ (Figure 5c), our results suggest that IHSs represent sites of cytolysis or injury followed by tissue regeneration. However, based on the technical limitations, further analyses, beyond the scope of this study, are necessary to validate this hypothesis in greater detail.

## DISCUSSION

In this study we employ Spatial Transcriptomics and snRNA-seq to explore host-parasite interactions during *P. berghei* liver stage development in the true tissue context. We uncover spatial elements that impact parasite growth and immune evasion, including tissue-wide and focal inflammatory responses, lipid homeostasis and liver zonation. Moreover, we evaluate the roles of myeloid and lymphoid immune cells along with other liver resident cells during malaria infection.

Recent advances in next generation sequencing have greatly enhanced our understanding of multiple stages of the *Plasmodium* life cycle, including liver stage development ^53–56^. However, until recently, spatial information of host-parasite interactions in liver tissue has been missing. While *Afriat* and colleagues described spatio-temporal interactions at the single-cell-level between zonated hepatocytes and *P. berghei* parasites, comprehensive investigations within the true tissue context have been lacking. This includes potential paracrine and endocrine interactions of infected hepatocytes and surrounding cells as well as other cell types.

Performing Spatial Transcriptomics with immunofluorescence staining of the intact parasites (UIS4) on the same infected tissue section, enabled us to associate transcriptional programs with parasite neighborhoods. We established correlations between gene expression involved in immune and lipid metabolism pathways near parasite neighborhoods at the late stages of infection. Moreover, we showed activation of various immune cell types during infection. Our analyses do not show a correlation of increased immune cell proportion near parasite neighborhoods, which suggests that immune cell activation may be uniformly distributed across the tissue and may effectively be evaded by successful parasites within the parenchyma.

Lipids are essential for *P. berghei* liver stage development and are scavenged from the host cells by the parasite ^57^. We speculate that the changes in lipid composition at the site of infection 38 hpi may exhibit anti-inflammatory effects by restricting recruitment of effector cells of the innate immune response to the site of infection ^58–60^. Our data indicate that there are no increased cell type proportions of immune effector cells in proximity to parasite positions at 38 hpi. Interestingly, our data show higher expression of *Fabp5* close to parasite locations, *Fabp5* is known to selectively enhance the activities of PPARß/∂ and PPARγ ^36^. It has previously been described that PPARs reduce inflammation by exhibiting anti-inflammatory potential ^61–64^. Thus, induced upregulation of expression of *Fabp5* may exhibit a lipid metabolism-dependent evasion strategy induced by the parasite. Meanwhile, *Insig1* expression increases with increased distance from the parasite. The absence of *Insig1* enhances lipid and cholesterol synthesis ^34^, potentially providing more lipid and cholesterol for the parasite in its proximity. Further, our analyses show upregulation of expression of the autophagy antagonist *Rheb* ^39,65^ in close proximity to parasite locations in the tissues. This observation suggests that downregulation of *Rheb* may assist *P. berghei* to evade elimination of host autophagy by limiting autophagosome formation ^39^.

Upon entering the liver, the parasite crosses the sinusoidal layer and continues to traverse multiple hepatocytes before invading a final hepatocyte, where it initiates replication ^82–84^. The reason for this traversal is still elusive ^66^ and detailed characterization of the interactions between traversed hepatocytes and immune cell responses remains a subject of investigation. Potentially, IFN-mediated immune responses are triggered by both traversed and infected cells, or result from paracrine crosstalk among infected, traversed, and neighboring immune and parenchymal cells. The high dose of sporozoites in our study may in part explain the global activation of previously reported upregulation of ISGs during infection progression ^15–17^.

We find that tissue-wide pro-inflammatory responses occur with a delay of 12 to 26 hours (at time points 24 and 38 post-challenge) in tissue sections from SGC mice. This delayed response is likely triggered by proteins from mosquito salivary glands and residual bacterial material in the saliva.

Future studies could further explore this finding using lower numbers of parasites, more in line with a natural infection and comparing different infection methods including increasing number of exposures to mosquito bites. It would also be interesting to compare how overall innate immune responses towards salivary gland components differ from responses to sporozoites beyond 38 hours post challenge.

Furthermore, we identified inflammatory hotspots (IHSs) with distinct tissue morphology, showing upregulated pro-inflammatory gene signatures nearby. This is supported by increased proportions of various immune cell types and cell surface markers near IHSs. These infiltrates, resembling responses to local inflammation, can have diverse cell compositions and effects on liver health, often involving immune response and regeneration ^23,24^. IHSs have been observed in viral diseases like rubella, COVID-19, and Epstein-Barr Virus, which affect the liver without causing significant liver disease, usually resulting in subclinical involvement and self-limitation ^22,23^. To our knowledge, these focal inflammatory infiltrates or IHSs have not previously been reported in the context of malaria. However, they might be of clinical relevance as it has been suggested that liver injury in clinical malaria is an overlooked phenomenon ^67^. We do not observe co-localization of IHSs with parasites stained with UIS4 antibodies, UIS4 has been ascribed a critical role in avoiding parasite elimination, suggesting the parasites we detect are still intact ^68^. Immune infiltration could be triggered by the parasite’s initial traversal through hepatocytes during early invasion, or by parasites that failed to successfully invade or develop early during the liver stage. Moreover, the location of the IHSs in close proximity to portal veins further highlights the importance of liver zonation for parasite survival, previously reported by Afriat et al. ^18^.

In our proposed model, malaria parasites not only resist pro-inflammatory host signals but may actively promote inflammation attenuation in their vicinity, thereby limiting the infiltration of effector immune cells. This evasion strategy involves the modulation of lipid homeostasis, including PPAR signaling and a limitation of autophagy. IHSs may form due to parasite traversal after entering the liver parenchyma or influence parasite elimination earlier than 12 hpi, affecting developmental success of the parasite (Figure 6). However, additional studies are necessary to fully characterize their role during malaria development in the liver.

**Figure 6.**
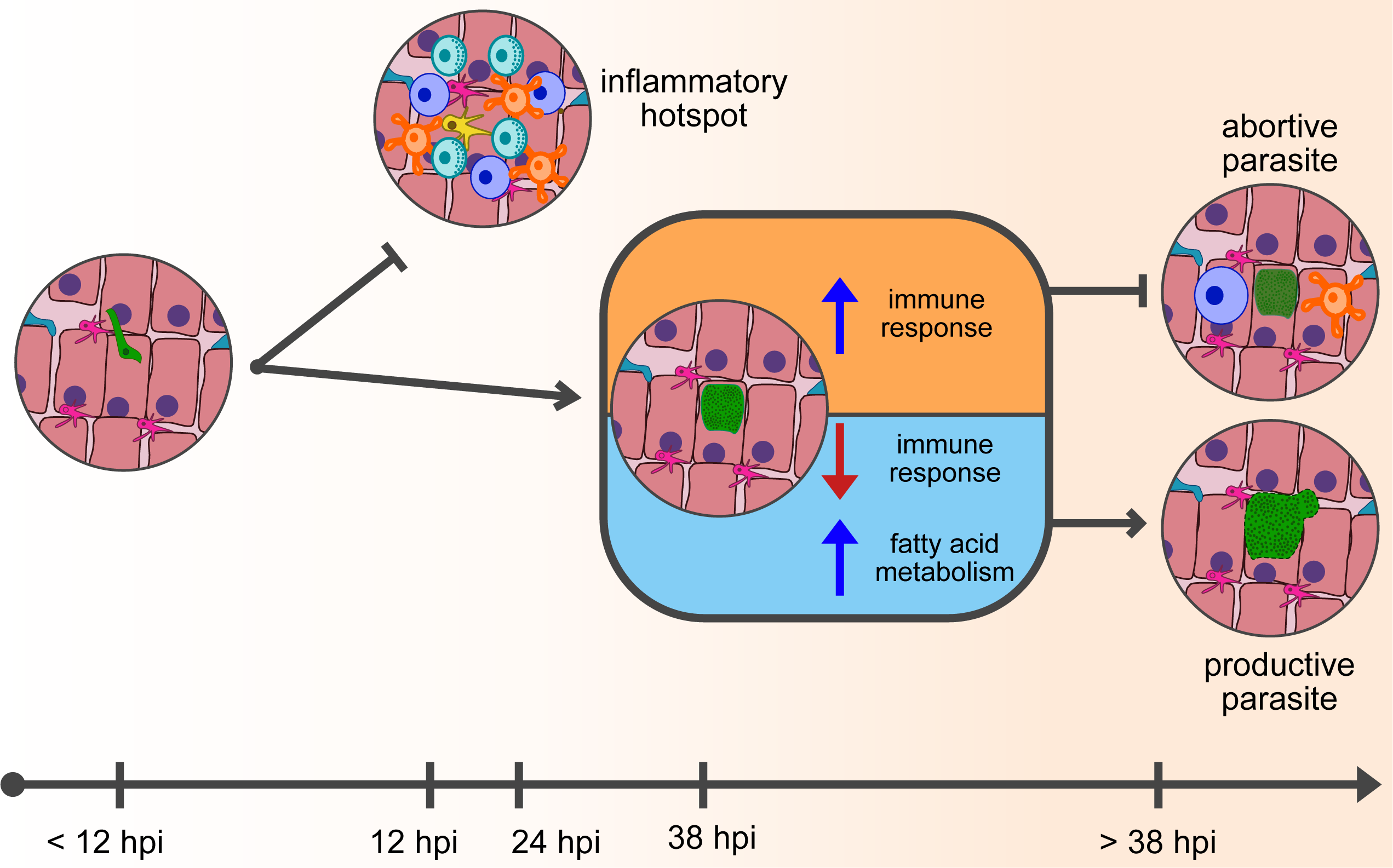
Proposed model for parasite clearance in the liver. Upon infection of the vertebrate host, the malaria parasite quickly enters the liver parenchyma and infects a hepatocyte (prior to the 12 hpi timepoint of this study). If the parasite, during this initial developmental phase, becomes exposed by the infected hepatocyte or surrounding non-parenchymal cells, the infection may result in clearance through immune cell infiltration, which would lead to the formation of an inflammatory hotspot. However, in the case that the infection persists towards late *P. berghei* liver-stage infection, the parasite down-regulates genes involved in a productive immune response, while deregulating fatty acid metabolism and autophagy within the vicinity of the infection site - thus, ensuring parasite proliferation. However, if the parasite fails to deregulate these pathways in its proximity during late liver-stage infection, the parasite will most likely be aborted.

In summary, our study provides a detailed spatiotemporal atlas of the host-parasite interplay during *Plasmodium* development in the liver, at the tissue level. Malaria eradication efforts require more extensive knowledge of the underlying biology in *de novo* immunization efforts. To this end, high-resolution spatial omics applications will be indispensable for understanding the coordination of immune priming in events of partial or full immunization. Future studies will also be necessary to broaden our understanding of the involvement of lipid metabolism, autophagy and IHSs reported in our study, which may provide novel avenues of combating malaria disease prior to reaching the symptomatic blood-stage infection.

## METHODS

### Ethical statement

The study was performed in strict accordance with the recommendations from the Guide for Care and Use of Laboratory Animals of the National Institutes of Health (NIH). The animal use was done in accordance with the National Institute of Allergy and Infectious Diseases Animal Care and Use Committees (NIAID ACUC), proposal LMVR 22.

### *P. berghei* infections and sample collection

Challenges with *Plasmodium berghei ANKA* (Anka 2.34) sporozoites or salivary gland lysate (uninfected *Anopeheles stephensi*) in female 8-9-week-old C57BL/6 mice were performed by tail-vein injection. First, *P. berghei* infected *A. gambiae* or *A. stephensi* salivary glands were collected 18-21 days post infection dissected to collect sufficient sporozoites for each challenge. The corresponding number of salivary glands were collected from non-infected mosquitoes for control challenges with salivary gland lysate. Sporozoites and lysate were pelleted by centrifugation, washed and stored in cold PBS, where the final concentration of sporozoites was determined. Sporozoites were diluted to reach a total number of 300,000 - 400,000 sporozoites for each infection. After tail vein injection, livers were collected after 12, 24 or 38 hours.

### Collection and preparation of liver samples

The livers were collected, and lobes were separated. Each lobe was segmented so cryosections would fit on the 6,200 x 6,400 µm areas of the Codelink-activated microscope or Visium slides and frozen in -30°C 2-Methylbutane (Merck, cat.no.: M32631-1L). For spatial experiments, the frozen liver samples were embedded in cryomolds (10×10 mm, TissueTek) filled with pre-chilled (4°C) OCT embedding matrix, frozen (CellPath, cat.no.: 00411243) and sectioned at 10 µm thickness with a cryostat (Cryostar NX70, ThermoFisher). Each subarray on the slide is covered with 1934 spots with a 100 µm diameter, containing millions of uniquely barcoded oligonucleotides with poly-T_20_ VN capture regions per spot (barcoded slides were manufactured by 10X Genomics Inc). The full protocol, including sequencing and computational analysis was performed for a total of 38 sections of which 23 were infected with *Plasmodium berghei* parasites and 15 challenged with mosquito salivary gland lysate. We analyzed 4 biological replicates for infected samples collected after 12h and 24h and 2 biological replicates for infected samples after 38h. For controls, we analyzed livers for 3, 3 and 2 biological replicates, respectively. Samples were selected based on sectioning and RNA quality.

### Immunofluorescence staining of spatial slides

We performed a modified version of the Spatial Transcriptomics workflow according to *Ståhl et al.* and *Vickovic et al.*, respectively ^69,70^. After placing the sections on the ST or Visium slides, they were fixed for 10 minutes using 4% formaldehyde in PBS. Then, they were dried with isopropanol and parasites were labeled using immunofluorescence as read-out. In short, after fixation, a blocking step using 5% Donkey-serum (Merck, cat.no: D9663-10ML) in PBS for 15 minutes was performed. Washing steps were performed using a 3 times concentrated SSC-buffer in deionized and RNAse-free water and RNAse Inhibitor (SUPERase•In™ RNase Inhibitor, Thermo Fisher Scientific, cat.no: AM2694), further referred to as blocking buffer. Staining of parasites was performed using an antibody against *Plasmodium berghei* UIS4 produced in goat (Nordic BioSite, cat.no: LS-C204260-400) in a concentration of 1:100 in 1:5 concentrated blocking buffer for 20 minutes at room temperature. The sections were washed and fluorescently labeled using a Donkey anti-Goat IgG (H+L) Highly Cross-Adsorbed Secondary Antibody, Alexa Fluor Plus 594 (Thermo Fisher Scientific, cat.no: A32758) at a concentration of 1:1000 in 1:5 concentrated blocking buffer for 20 minutes at room temperature and in the dark. The slides were washed and DNA was stained using 1:1000 concentrated DAPI solution (Thermo Fisher Scientific, cat.no:62248) for 5 minutes at room temperature and in the dark. Then, slides were mounted with 85% glycerol (Merck Millipore, cat.no.: 8187091000) including RNAse Inhibitor (SUPERase•In™ RNase Inhibitor, Thermo Fisher Scientific, cat.no: AM2694) and covered with a coverslip. Images were acquired at 20x magnification, using the Zeiss AxioImager 2Z microscope and the Metafer Slide Scanning System (Metasystems).

### Histological staining and annotations

After immunofluorescence staining, a histological staining with Mayer’s hematoxylin (Dako, cat.no.: S330930-2) followed by Eosin (Sigma-Aldrich, cat.no.: HT110216-500ML), diluted in Tris/acetic acid (pH 6.0) was performed. The stained sections were mounted with 85% glycerol (Merck Millipore, cat.no.: 8187091000) and covered with a coverslip. Bright field images were acquired at 20x magnification, using Zeiss AxioImager 2Z microscope and the Metafer Slide Scanning System (Metasystems). The liver images were assessed by an expert liver histologist (NVH) who annotated the portal (PV) and central veins (CV), based on the presence of bile ducts and portal vein mesenchyme (PV) or lack thereof (CV). When the quality of the sample did not allow for annotation, “ambiguous vein” was reported. Moreover, regions of apparent cell infiltration were annotated based on increased nuclear signal.

### Permeabilization, cDNA synthesis, tissue removal and probe release

Next, the slides were put in slide cassettes to enable separated on-array reactions in each chamber as described previously ^70^. Each tissue section was pre-permeabilized using Collagenase I for 20 minutes at 37°C. Permeabilization was performed using 0.1% pepsin in 0.1 M HCl for 10 minutes at 37°C. cDNA synthesis was performed overnight at 42°C. Tissue removal from the arrays prior to probe release was performed using Proteinase K in PKD buffer at a 1:7 ratio at 56°C for 1 hour. Lastly, the surface probes were released and cDNA library preparation followed by sequencing was performed.

### cDNA library preparation and sequencing

Released mRNA-DNA hybrids were further processed to generate cDNA libraries for sequencing. In short, the 2^nd^ strand synthesis, cDNA purification, in vitro transcription, amplified RNA purification, adapter ligation, and post-ligation purification, were done using an automated MBS 8000+ system ^71^. To determine the number of PCR cycles needed for optimal indexing conditions, a qPCR was performed. After determination of the optimal cycle number for each sample, the remaining cDNA was indexed, amplified and purified ^72^. The average length of the indexed cDNA libraries was determined with a 2100 Bioanalyzer using the Bioanalyzer High Sensitivity DNA kit (Agilent, cat.no.:5067-4626), concentrations were measured using a Qubit dsDNA HS Assay Kit (Thermofisher, cat.no:Q32851) and libraries were diluted to 4nM. Paired-end sequencing was performed on the Illumina NextSeq500 (v2.5 flow cell) or NextSeq2000 platform (p2 or p3 flow cell), resulting in the generation of 80 to 150 million raw reads per sample. To assess the quality of the reads FastQC (v 0.11.8) reports were generated for all samples.

### Spot visualization and image alignment

The staining, visualization and imaging acquisition of spots printed on the ST slides were performed. Briefly, spots were hybridized with fluorescently labeled probes for staining and subsequently imaged on the Metafer Slide Scanning system (Metasystems). The previously obtained brightfield of the tissue slides and the fluorescent spot images were then loaded in the web-based ST Spot Detector tool ^73^. Using the tool, the images were aligned and the spots under the tissue were recognized by the built-in recognition tool. Spots under the tissue were then slightly adjusted and extracted.

### Visium experiments

Spatial experiments with increased resolution were carried out using the 10X Visium Spatial Technology (10X Genomics, cat.no: 1000187) according to a slightly modified version of the protocol provided by 10X Visium. In brief, immunofluorescent staining of *P. berghei* parasites using an anti-UIS4 antibody and DNA using DAPI was performed as described above. After fluorescent imaging, Hematoxylin & Eosin (H&E) staining and brightfield imaging, the tissue was permeabilized for 30 minutes using the permeabilization buffer provided by the reaction kit. Then cDNA synthesis, template-switching and second strand synthesis were performed according to the protocol. Library generation was performed by amplification and purification of resulting products from the previous steps. Fragment traces were determined with a 2100 Bioanalyzer using the Bioanalyzer High Sensitivity DNA kit (Agilent, cat.no.:5067-4626), concentrations were measured using a Qubit dsDNA HS Assay Kit (Thermofisher, cat.no: Q32851) and libraries were diluted to 2nM and pooled for sequencing. Sequencing was performed using a NextSeq2000 (p2 or p3 flow cell) instrument resulting in approximately 80 million reads per sample.

### Single-nuclei RNA-sequencing (snRNA-seq)

Nuclei were isolated from snap frozen liver tissue with a sucrose gradient as previously described ^74^. Briefly, frozen liver tissue was homogenized using the Kimble Dounce grinder set to 1 ml in the homogenization buffer with RNAse inhibitors. Homogenized tissue was then subjected to density gradient (29% cushion – Optiprep) ultracentrifugation (7700rpm, 4°C, 30 mins). Nuclei were resuspended and 2 biological replicates of each condition were pooled before nuclei were stained using DAPI. Intact nuclei were FACS-purified from remaining debris. A total of 60000 nuclei were sorted into BSA coated tubes. The sorted nuclei were pelleted by centrifugation for 3 mins at 400g and 5 mins at 600g, sequentially. Nuclei were then resuspended in PBS with 0.04% BSA at ∼1000 nuclei/µl. Nuclei suspensions (target recovery of 20000 nuclei) were loaded on a GemCode Single-Cell Instrument (10x Genomics, Pleasanton, CA, USA) to generate single-cell Gel Bead-in-Emulsions (GEMs). Single-cell RNA-Seq libraries were prepared using GemCode Single-Cell 3□Gel Bead and Library Kit (10x Genomics, V2 and V3 technology) according to the manufacturer’s instructions. Briefly, GEM-RT was performed in a 96-Deep Well Reaction Module: 55°C for 45 min, 85°C for 5 min; end at 4°C. After RT, GEMs were broken down and the cDNA was cleaned up with DynaBeads MyOne Silane Beads (Thermo Fisher Scientific, 37002D) and SPRIselect Reagent Kit (SPRI; Beckman Coulter; B23318). cDNA was amplified with 96-Deep Well Reaction Module: 98°C for 3 min; cycled 12 times: 98°C for 15s, 67°C for 20 s, and 72°C for 1 min; 72°C for 1 min; end at 4°C. Amplified cDNA product was cleaned up with SPRIselect Reagent Kit prior to enzymatic fragmentation. Indexed sequencing libraries were generated using the reagents in the GemCode Single-Cell 3□ Library Kit with the following intermediates: (1) end repair; (2) A-tailing; (3) adapter ligation; (4) post-ligation SPRIselect cleanup and (5) sample index PCR. Pre-fragmentation and post-sample index PCR samples were analyzed using the Agilent 2100 Bioanalyzer.

snRNA-seq libraries were pooled in equal ratios and loaded on a S4 lane Illumina NovaSeq 6000, resulting in 2500 - 3000 million read-pairs. Sequencing was performed at the National Genomics Platform (NGI) in Stockholm, Sweden. Spatial (Spatial Transcriptomics, Visium) and snRNA-seq data were aligned to a combined custom reference genome combining *Mus musculus* (GRCm38.101) and *Plasmodium berghei* (PlasmoDB-48_PbergheiANKA) using stpipeline ^75^(v.1.8.1) and STAR (v.2.6.1e), spaceranger (v.2.0.0) or cellranger (v.3.0.0), respectively.

### Immunofluorescence staining of inflammatory hotspots

We performed IF staining of *P. berghei* infected and control (salivary gland lysate challenged) tissues after 12, 24 and 38 hpi. For each experiment, three consecutive tissue sections of the same tissues utilized for spatial as well as single nuclei experiments were placed on spatially separated positions of a Super frost slide (VWR, cat.no: 631-0108). After placement, the tissue was fixed using pre-cooled methanol and incubated for 15 minutes at -20°C. Tissue sections were permeabilized using 0.2% TritonX-100 (Sigma, cat.no: T8787) in PBS for 5 minutes and blocked for 15 minutes using 5% donkey-serum in PBS. After blocking, mouse specific primary antibodies were applied in different combinations across the three sections. These included i) <u>10 µg/ml</u> monoclonal CD4 (Thermo Fisher, cat.no: MA1-146, clone GK1.5), and <u>10 g/ml</u> monoclonal CD8 (Thermo Fisher Scientific, cat.no: MA5-29682, clone 208), ii) <u>1:100 diluted</u> monoclonal F4/80 (Thermo Fisher Scientific, cat.no: MA5-16624, clone CI:A3-1), and <u>2 µg/ml</u> monoclonal CD27 (Thermo Fisher Scientific, cat.no: MA5-29671, clone 12) and iii) <u>10 µg/ml</u> monoclonal CD11b (Thermo Fisher Scientific; cat.no: 53-0112-82, clone M1/70) and <u>5 µg/ml</u> monoclonal CD11c (Thermo Fisher Scientific, cat.no: 42-0114-82, clone N418). All antibodies were incubated with the tissue for 60 minutes at room temperature. Tissue sections were washed three times with PBS and corresponding secondary antibodies were applied. These included i) Donkey anti-Rat IgG (H+L) Highly Cross-Adsorbed Secondary Antibody, Alexa FluorTM 488, InvitrogenTM (cat.no: A21208) ii) Donkey anti-Rabbit IgG (H+L) Highly Cross-Adsorbed Secondary Antibody, Alexa Fluor™ 555 (cat.no: A-31572), iii) Donkey anti-Rat IgG (H+L) Highly Cross-Adsorbed Secondary Antibody, Alexa FluorTM 647 (cat.no: A78947) and iv) Donkey anti-Rabbit IgG (H+L) Highly Cross-Adsorbed Secondary Antibody, Alexa FluorTM Plus 647 (cat.no: A32795). All antibodies were incubated with the tissue for 30 minutes at room temperature. Tissue sections were washed three times with PBS and DNA was stained using (1 µg/ml) DAPI (Thermo Fisher Scientific, cat.no: 62248) for 5 minutes at room temperature. Tissue sections were mounted using Diamond antifade mounting medium (Thermo Fisher Scientific, cat.no: S36972) and imaged. To select inflammatory hotspots which occur in all three consecutive sections, a tiled scan of the DNA counterstain was performed at 20X magnification. Selected hotspots were then imaged at 40X magnification using the same settings across each tissue section. Imaging analysis was performed using ImageJ, were brightness and contrast were adjusted for visualization purposes and composite creation.

### Computational analysis

#### Filtering, normalization, integration, dimensionality reduction and unsupervised clustering

Main computational analysis of spatial read-count matrices (ST and Visium) was performed using the STUtility package (v 0.1.0) ^76^ in R (v 4.0.5). The complete R workflow can be assessed and reproduced in R markdown (see code availability section). Analysis of snRNA-seq data was in large parts performed using the Seurat package (v 4.1.1). For ST and Visium data only protein coding genes were considered for analysis and genes of the major urinary protein (Mup) family were filtered due to the large differences in expression between individual mice ^18,77^. Gene esxpression was normalized, accounting for differences sequencing depth and circadian effects caused by the dissection time point. Subsequently, normalized expression data was scaled and highly variable genes were selected using the SCTransform function in Seurat. All samples, biological replicates and dissection time points were further corrected for batch effects using the harmony package (v.0.1.0) ^78^. Thereafter, the first 20 harmony vectors were subjected to shared-nearest-neighbor (SNN) inspired graph-based clustering via the *“FindNeighbors*” and “*FindClusters*” functions. For modularity optimization, the Louvain algorithm was used and clustering was performed at a resolution of 0.35 for clustering granularity.

#### Visualization and spatial annotation of clusters

To visualize the clusters in low-dimensional space for snRNA-seq and spatial data as well as the spot coordinates under the tissue for spatial data, non-linear dimensionality reduction was performed using UMAP. Visualization and annotation of identified clusters in UMAP space (snRNA-seq, ST, Visium) on spot coordinates as well as superimposed on the H&E images (ST, Visium) was performed using the Seurat and STUtility package.

#### Differential gene expression analysis and gene modules in space

To investigate changes in gene expression between selected groups, differential gene expression analysis (DGEA) was performed. Groups for comparison were selected in a supervised (tested conditions) or unsupervised fashion (clustering). Then the FindAllMarkers function of the Seurat package was employed to identify all differentially expressed genes (DEGs) between all investigated groups, including genes with a logarithmic fold change above 0.25. Only DEGs below an adjusted p-value of 0.05 were considered for further downstream analysis. To investigate differentially expressed genes between two groups only, the FindMarkers function of the Seurat package was employed using the same thresholds as described. In both cases a Wilcoxon-rank sum test was performed to identify differentially expressed genes.

#### Functional enrichment analysis

Functional enrichment of genes of interest was performed using the grpofiler2 package (v.1.0). The algorithm defined in the “gost” function takes a list of genes and associates them with known functional information sources, establishing statistically significant enriched terms. This package is able to take data from mouse and several other organisms into account to perform the analysis, but lacks data of *P. berghei* or other *Plasmodium* species. Therefore, functional enrichment analysis was only performed for *Mus musculus* genes. We investigated functional enrichment from the KEGG and Gene Ontology (GO) database sources and significance was adjusted using g:SCS (Set Counts and Sizes) ^79^. Visualization was performed for the most highly enriched terms and enrichment scores are represented as the negative log10 algorithm of the corrected p-value.

#### Cluster Interaction Analysis

To approximate how expression-based clusters interacted in the tissue space, a simple interaction analysis was carried out as described in detail previously ^19^. Briefly, the cluster identity or the four nearest-neighboring spots within a distance threshold were registered, to ensure spots located in the actual physical neighborhood were included in the count, as this assumption might not hold for spots at the edge of the tissue. A binomial test was performed to test for significant over (or under) representation (Cluster interactions) and resulting values were visualized in a heatmap and grouped hierarchically, using complete linkage clustering, in the seaborn package (v.0.12.2) in python (v 2.7.18). Based on the fact that clusters vary considerably in size, a random permutation of cluster positions was performed to investigate which interactions are likely to be occurring by chance.

#### Features as a function of distance

To investigate the relationship between features of interest (gene expression, proportion values) and the distance to a structure of interest (vasculature, parasites, inflammation hotspots) in the tissue sections, the values of the features of interest were modeled as a function of the distance as previously described ^19^. In short, brightfield or fluorescence images were used to create a mask for each structure of interest. As the position of the capture locations relate to the pixel coordinates in the H&E images, the created masks were used to computationally measure the distance from each spot to each selected structure. The distance to a selected structure was defined as the minimal euclidean distance from the center of each spot to any pixel of the union of all masks.

#### Expression-by-distance analysis and distance-based correlation analysis

After determining distances of spots (capture locations), the distance to each structure of interest was associated with each spot and used for downstream analyses, and visualization was adapted using similar to previously reported visualization approaches ^19^.

To investigate the relationship between a structure of interest and gene expression in its neighborhood across sections, Pearson correlations between the distance to the structure and expression values of each gene in the spatial gene expression data were performed. Spots within a threshold of 400 - 800 µm from the region of interest were selected. This was based on the size of the region of interest, with a threshold of 400 µm for smaller structures (e.g. parasites) and a threshold of 800 µm for larger structures (e.g. inflammation hotspots). After calculating correlations between distance and gene expression values, only adjusted (Bonferroni correction) significant correlations were selected (p <0.05) further.

Visualization of spatial relationships was carried out by plotting expression of correlated genes defined as *Y* over the distance to the structure of interest defined as *X*. To better capture trends of each relationship, loess smoothing X ∼ Y was applied to the data, similar as previously described ^19^. To better compare differences between different investigated conditions in some cases, the data were transformed to center around 0 for each condition of interest. This was performed by subtracting the fitted value of the loess regression at the minimal distance from each value in the expression data, maintaining the difference in expression Δ*Y* along the distance axis *X*. The ribbons around the smoothed curve represent the standard error (SE) as given by the loess algorithm.

#### Expression-based classification

Expression-based classification was performed for central and portal veins as previously described ^19^ using the hepaquery package (v.0.1). In brief, neighborhood expression profiles were created as described above (features as a function of distance) setting a threshold of 142 pixels, which refers to 400 µm and represents the longest distance between adjacent spot centers in the same row on an ST slide. After the formation of the neighborhoods, their associated weighted profiles for each gene were assembled. For each neighborhood, expression profile class label predictions were performed employing a logistic regression using the *LogisticRegression* class from *sklearn’s* (v 0.23.1) *linear_model* module in python. A *l2* penalty was used (regularization strength 1), the number of max iterations was set to 1000, default values were used for all other parameters. Performance validations were carried out using multiple levels of cross-validation as previously described ^19^. To prevent overfitting in the applied model due to the limited amount of structures, a reduced set of genes was used for the classification ^19^.

#### Single cell analysis and cell type annotation

The raw sequencing data files (.bcl files) were demultiplexed into FASTQ files using *cellranger mkfastq* (Cell Ranger v3.1.0, 10x Genomics) with default parameters. The demultiplexed reads were aligned to a custom genome of reference using the CellRanger (10x Genomics) pipeline. The genome of reference was created by combining the genomes of *Mus musculus* (GRCm38.101) and *Plasmodium berghei* (PlasmoDB-48_PbergheiANKA). This resulted in an expression matrix for each of the six sequenced samples (12, 24 and 38 hours infected and salivary gland control liver samples) which were individually analyzed. The quality control and clustering steps were performed using the *seurat* package (v.4.3.0) and following the standard workflow. The quality control pipeline involved (i) removing genes that were detected in fewer than 10 cells, (ii) filtering out cells with less than 200 genes and more than 5000 genes, (iii) excluding cells with over 15% mitochondrial transcripts and, (iv) discarding all mitochondrial and ribosomal genes from the expression matrix.

Doublets in the data were removed using *DoubletFinder* (v 2.0.3) with a pk of 0.005, 0.22, 0.24, 0.28 or 0.3, depending on the sample. Following this, the data was normalized and scaled using *SCTransform* (v0.3.5) with default parameters. The high variable genes needed to perform a principal component analysis (PCA) were identified using the *FindVariableFeatures* with the ‘vst’ method.

After initial filtering and doublet removal, gene expression of all investigated conditions were integrated using the *harmony* package (v0.1.0), defining the sample origin as a grouping variable. The *FindNeighbors* and *FindClusters* functions were used for clustering, and the Louvain algorithm was employed to cluster the cells with a resolution of 0.3 granularity.

Subsequently, cell type annotations were performed on the integrated data using a twofold strategy. First, an automatic cell type prediction was performed using scmap (v.1.16.0). For this, the top 500 most informative features for annotation were calculated using the *selectFeatures* function and the steady-state annotated mouse liver data set ‘Mouse StSt’, generated by *Guilliams et al.* ^80^ as a reference. Then, the scmap-cell pipeline was used to project the cell-type labels from the reference dataset onto our data. Following automatic annotation, a manual annotation step based on canonical marker genes was carried out which involved confirming and refining the obtained results from the automated annotation.

To calculate cell type proportions of immune cells (T and NK cells, B cells, monocytes and DCs and Kupffer cells) across conditions, the annotated cell data for infected samples (12, 24, and 38 hpi) and for control samples (12, 24 and 38 SGC) were each analyzed across infection time points. The average number of cell types of interest for the infected or control groups were calculated, and their proportions were obtained by dividing the cell type count by the total number of cells of the infected samples or the control samples, respectively. To assess the significance of differences between the three infected (12, 24, and 38 hpi) and the three control samples (12, 24 and 38 SGC), a two-sample t-test was performed using base R (v4.2.2).

#### Single cell data integration (*stereoscope*)

We integrated our annotated snRNA-seq data using *stereoscope* (v.0.3.1), a probabilistic method designed for spatial mapping of cell types ^81^. In short, *stereoscope* models both single cell and spatial data as negative binomial distributed, learns the cell type specific parameters and deconvolves the gene expression in each spot into proportion values associated with the respective cell type.

Stereoscope was run with 50,000 epochs and a batch size of 2048 for both sn and st modalities using subsetted snRNA-seq data and a list of highly variable genes. The annotated snRNA-seq expression matrix was subsetted to include a minimum of 25 and maximum of 250 cells per cell type, which were selected randomly. The list of highly variable genes was extracted from snRNA-seq data using Seurat (v.4.3.0) by first normalizing the data (*NormalizeData*, default parameters) and then identifying the highly variable genes (*FindVariableFeatures*, *selection.method = "vst", features = 5000*).

## Author contributions

**JA**, **JL and JVR** conceived and supervised the study. **FH** performed spatial transcriptomics experiments and analyzed the data. **FH, MUI, CZ** and **BV** extracted single nuclei and generated libraries. **FH** and **MUI** analyzed snRNA-seq data. **JVR, FH** and **TP** infected mice and dissected livers. **JVR** and **TP** dissected mosquitos. **NVH** performed histological annotations of liver tissues. **ES** and **FH** researched genes for gene set selections in expression-by-distance to the parasite analysis. **SS** and **MH** performed the stereoscope deconvolution and **FH** analyzed the data. **ES** performed immunofluorescence staining and analysis of inflammatory hotspots **ERA**, **CLS**, **JL** and **JVR** provided resources for the study. **FH**, **MUI** and **JA** wrote the manuscript. All authors provided feedback and edited the manuscript.

## Acknowledgments

We would like to thank the National Genomics Infrastructure in Stockholm funded by SciLifeLab, the Knut and Alice Wallenberg Foundation and the Swedish Research Council, as well as SNIC/Uppsala Multidisciplinary Center for Advanced Computational Science for assistance with massively parallel sequencing and access to the UPPMAX computational infrastructure. Parts of the computations were performed using resources provided by SNIC through Uppsala Multidisciplinary Center for Advanced Computational Science (UPPMAX). We thank Christian Gnann for feedback on the design of the antibody-based tissue-staining. We thank Amparo Roig Adam for feedback on the automated annotation of cell types. This study was generously funded by grants from: the Swedish Society for Medical Research (SSMF Establishment Grant), STINT, the Jaenssen Foundation and the Swedish Research Council (VR 2021-05057) to JA; the Sven and Lily Lawski foundation to FH; the NIH Distinguished Scholars Program and the Intramural Research Program of the Division of Intramural Research (AI001250-01), National Institute of Allergy and Infectious Diseases (NIAID), NIH to JVR; Karolinska Institutet (2-195/2021) and the Swedish Research Council (VR 2019-01350) to ERA and NVH; Marie Skłodowska-Curie Individual Fellowship (101027317, ‘MACtivate’) to CZ.

## Declaration of Disclosure

The authors do not declare any conflicts of interest. **SS**, **MH**, and **JL** are scientific advisors to 10x Genomics Inc, which holds IP rights to the ST technology.

## Data availability

Data will be made available on GeneExpression Omnibus and will be deposited in a zenodo repository upon publication.

## Code availability

Code to reproduce the analysis will be made available on Github (https://github.com/ANKARKLEVLAB) upon publication. Instructions for the installation and the workflow of the hepaquery package are already available at https://github.com/almaan/ST-mLiver.

## Notes

### Competing Interest Statement

The authors have declared no competing interest.

