## Supplementary figures for "Host-Pathogen Interactions in the *Plasmodium*-Infected Mouse Liver at Spatial and Single-Cell Resolution"

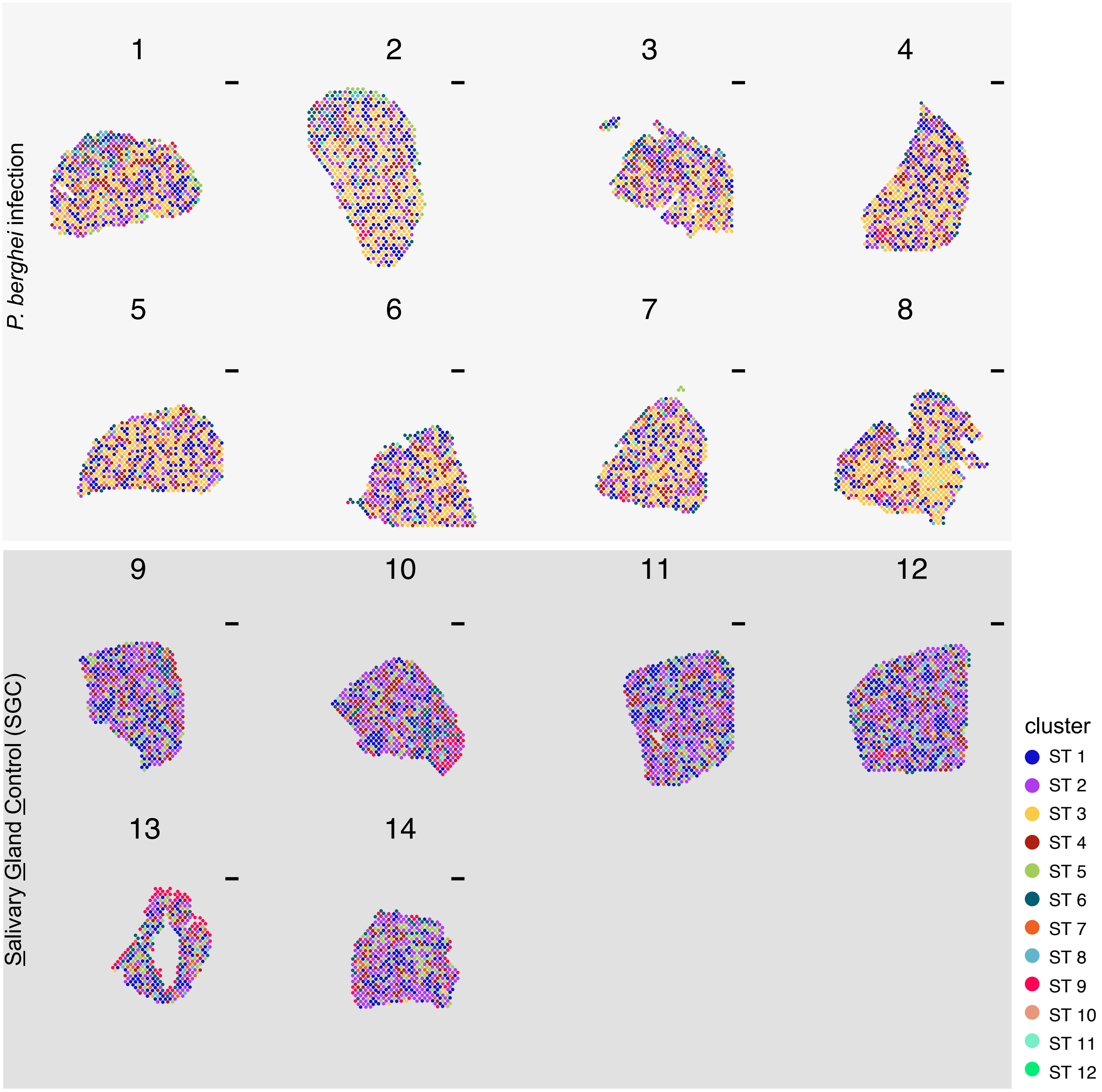


**Supplementary figure 1: Distribution of identified clusters across tissue sections at 12 hpi and respective control sections.** Predominant gene expression profiles according to clustering analysis across tissue sections 12 hpi (top, light grey) or 12h after salivary gland challenge (SGC) (bottom, dark grey). Different clusters (ST1 - ST12) are colored according to the legend (bottom right).

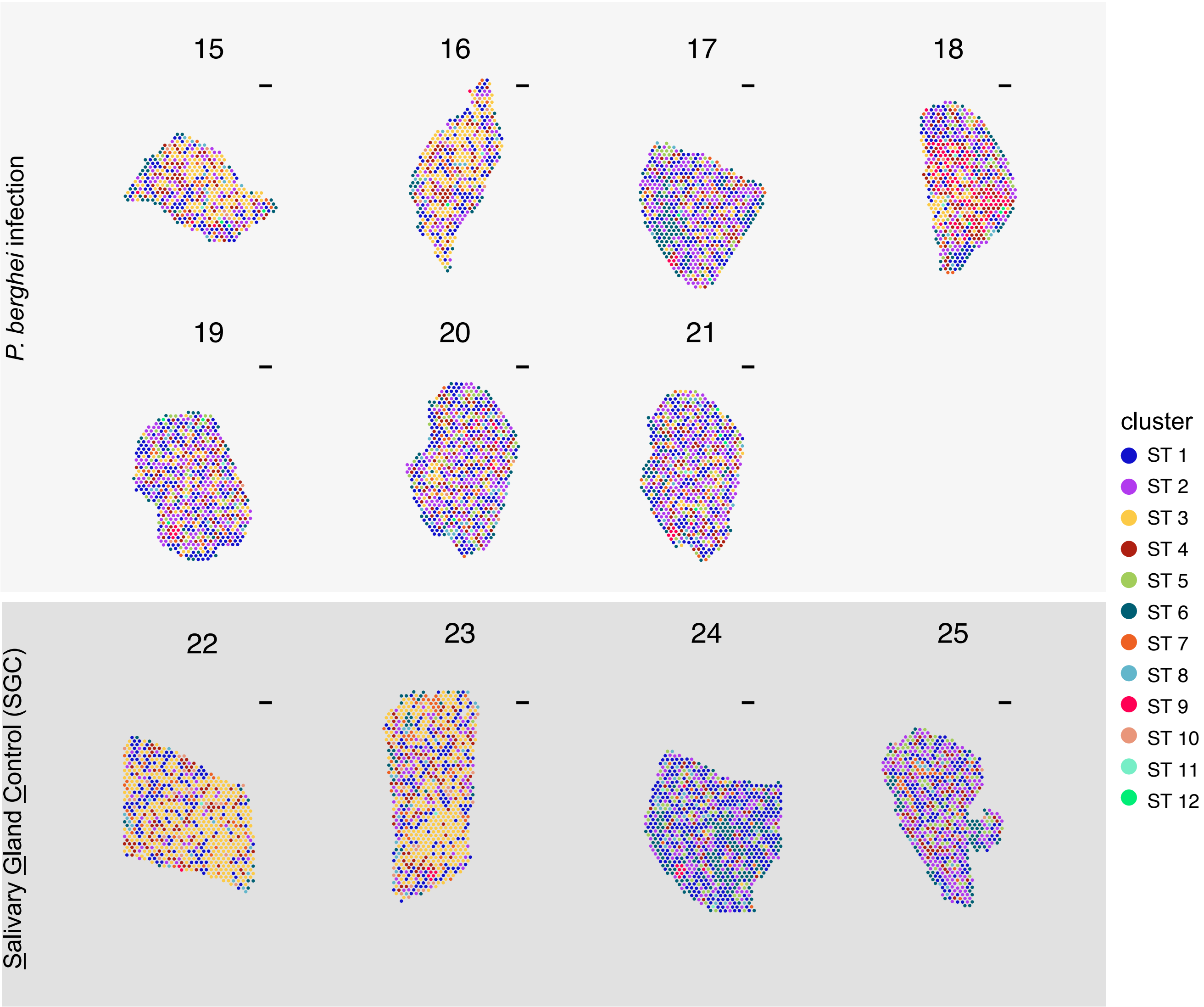


**Supplementary figure 2: Distribution of identified clusters across tissue sections at 24 hpi and respective control sections.** Predominant gene expression profiles according to clustering analysis across tissue sections 24 hpi (top, light grey) or 24h after salivary gland challenge (SGC) (bottom, dark grey). Different clusters (ST1 - ST12) are colored according to the legend (bottom right).

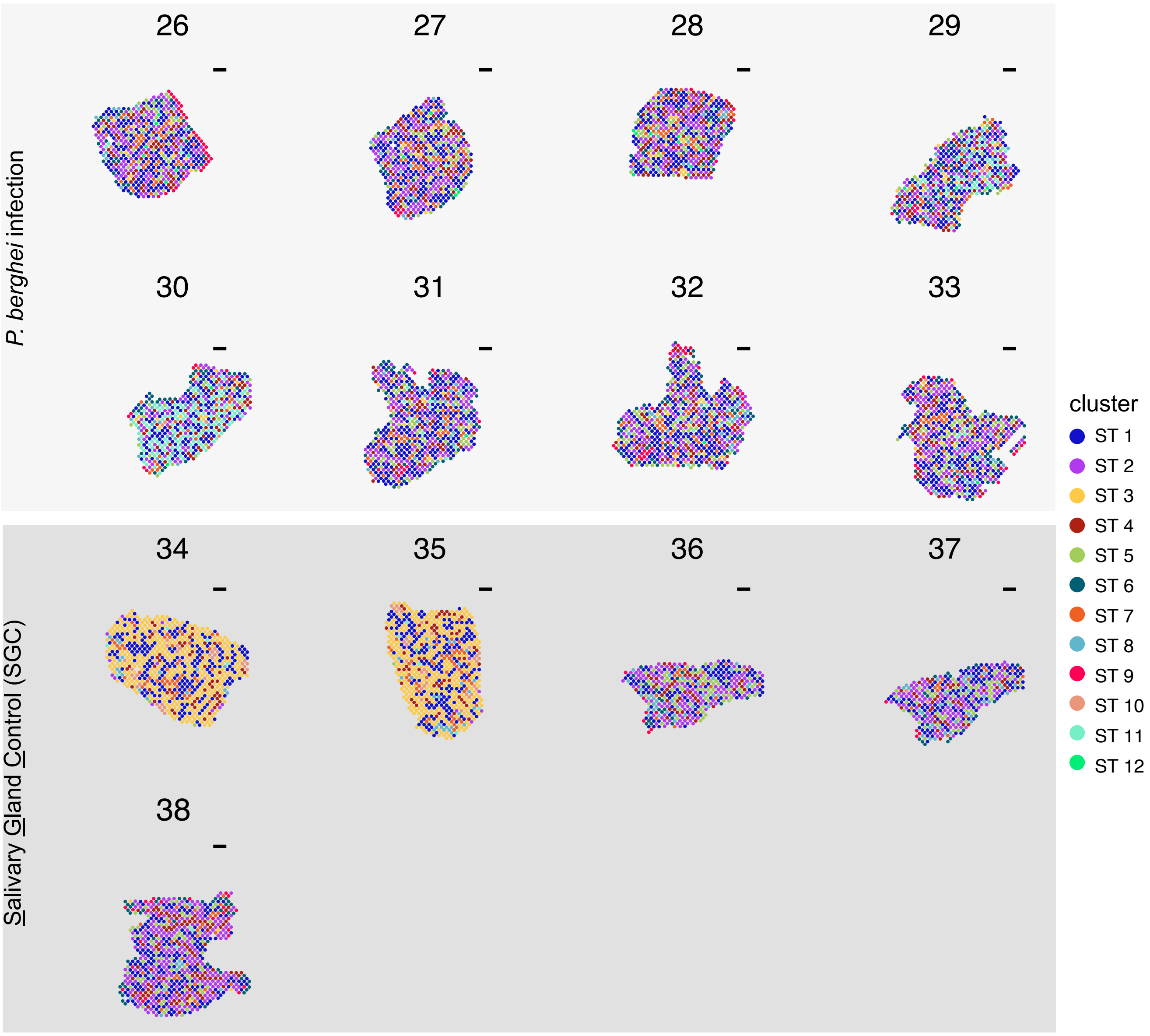


**Supplementary figure 3: Distribution of identified clusters across tissue sections at 38 hpi and respective control sections.** Predominant gene expression profiles according to clustering analysis across tissue sections 38 hpi (top, light grey) or 38h after salivary gland challenge (SGC) (bottom, dark grey). Different clusters (ST1 - ST12) are colored according to the legend (bottom right).

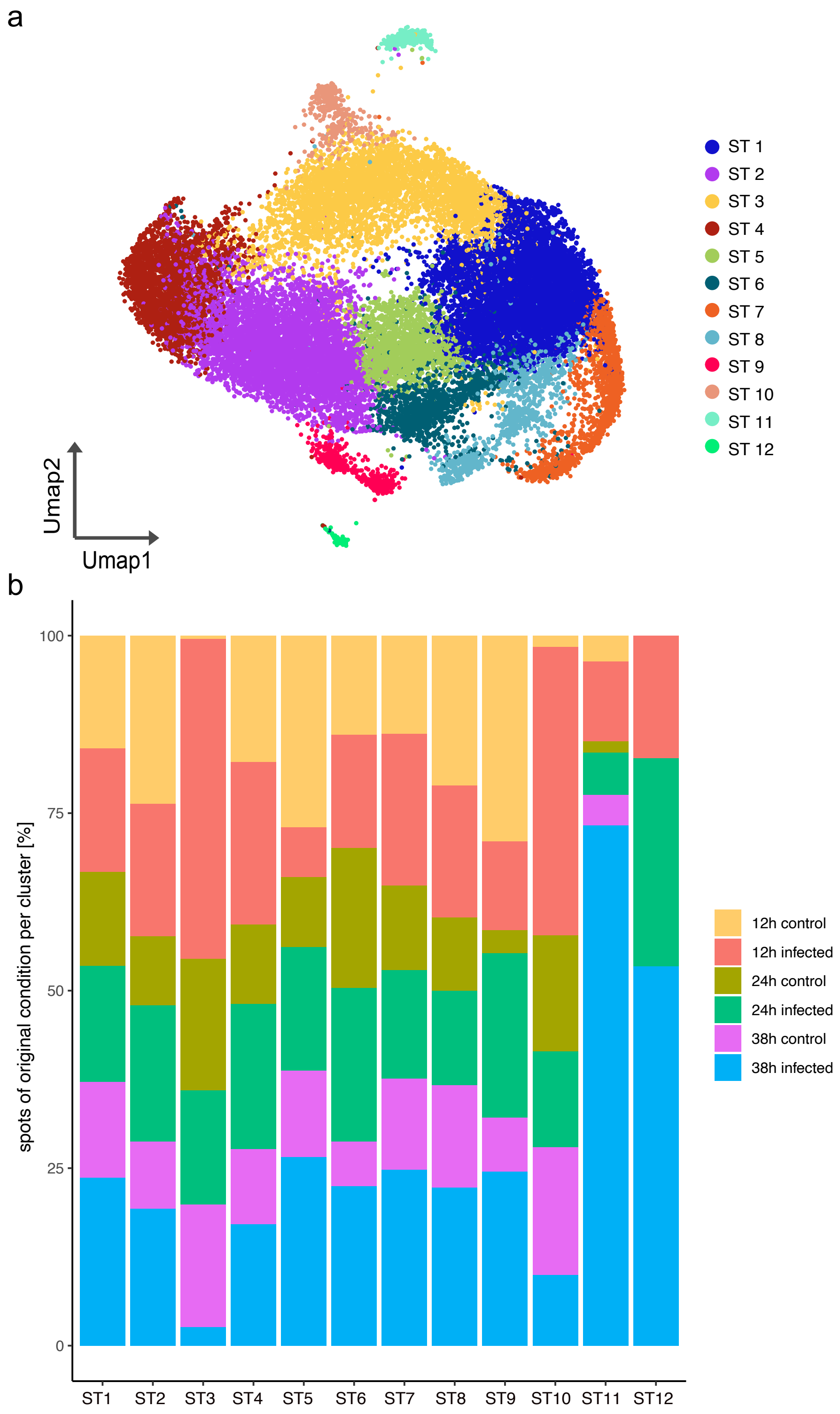


**Supplementary figure 4: Proportions of identified spatial clusters (ST1 - ST12) across infection conditions. a** UMAP emedding of clusters identified by unsupervised clustering **b** Percentages of spots of original conditions (12 - 38 hpi and respective controls) across all identified clusters from Spatial Transcriptomics (ST) analysis. Original conditions are indicated by color (12h control = yellow, 12h infected = red, 24h cotrol = olive, 24h infected = green), 38h control = pink, and 38 infected = blue).

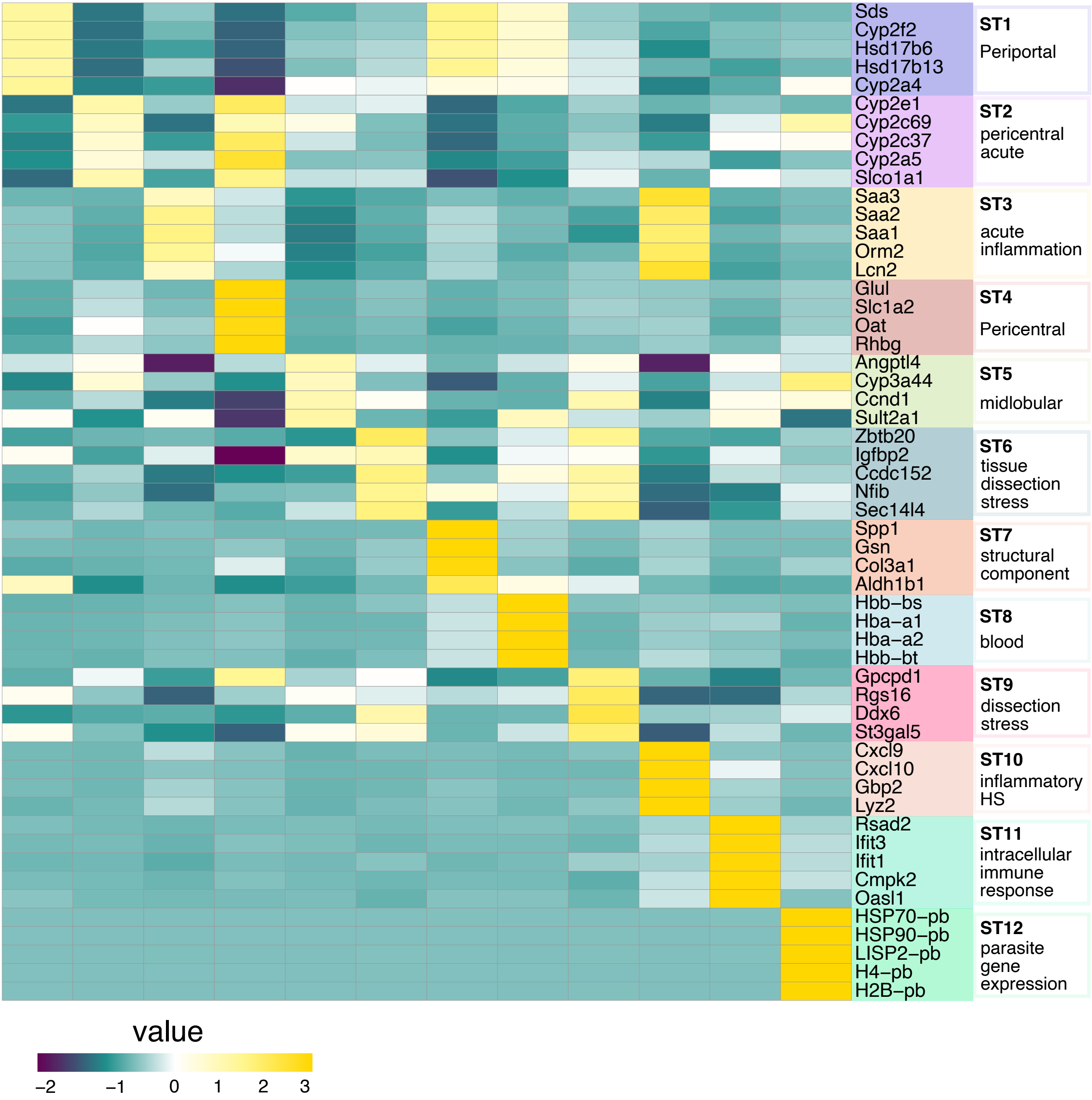


**Supplementary figure 5: Differential gene expression between Spatial Transcriptomics (ST) clusters.** Average gene expression of the top differentially expressed genes between identified spatial clusters. Spatial clusters are denoted by color starting with ST1 on the top and ST12 at the bottom. Function based annotations of clusters are shown on to the right of gene panels.

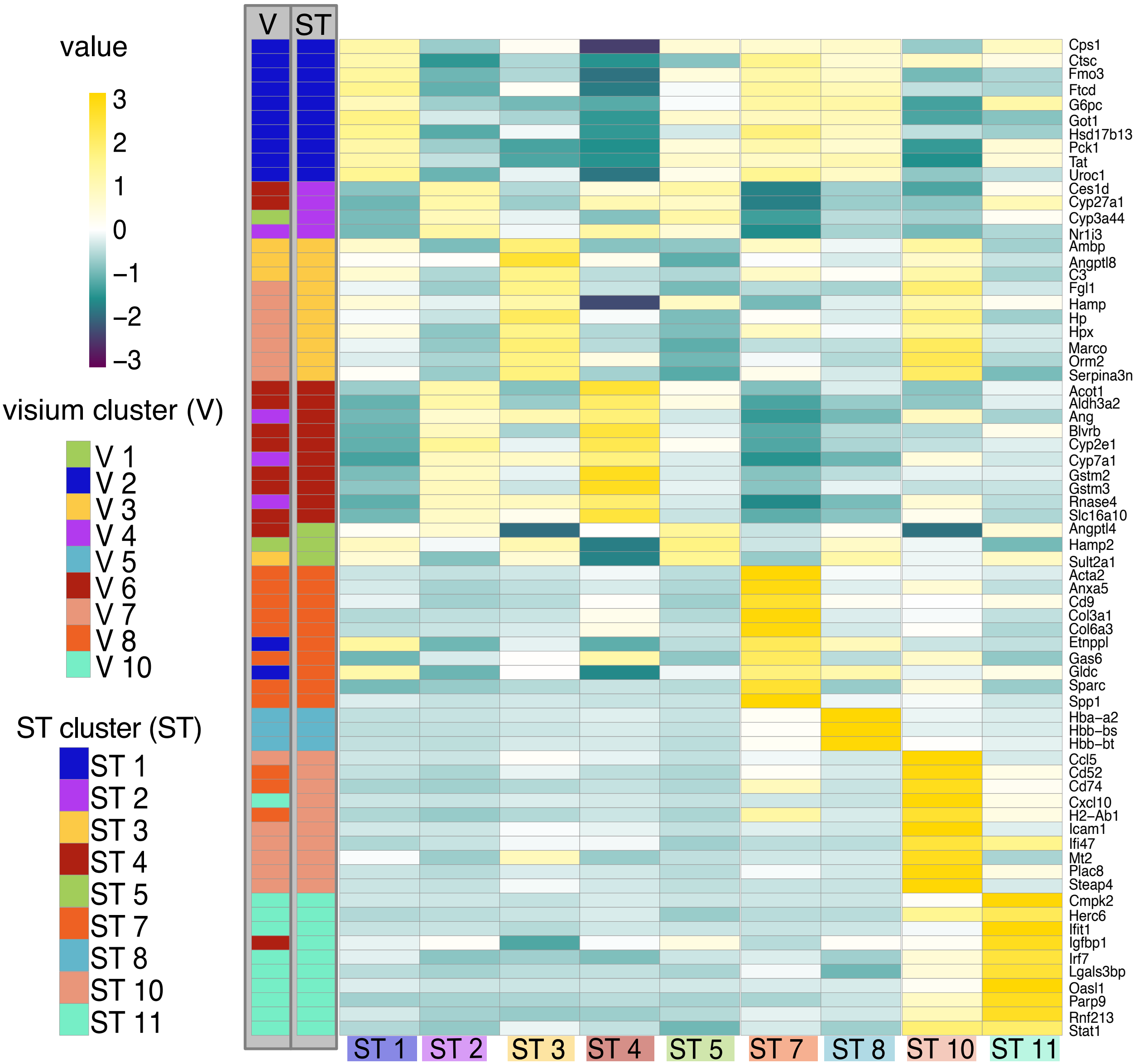


**Supplementary figure 6: Top shared differentially expressed genes between spatial technologies (ST) and Visium (V).** Differentially expressed genes between both spatial technologies were intersected and the average expression of hits were visualized in a heatmap using ST spatial data. Visium and ST clusters are depicted as colors as indicated by legends (left). Cluster associations for DEGs on the right are shown in the ST or V column on the left. Gene expression values are shown as a colorgradient from low expression (dark purple) to high expression (yellow).

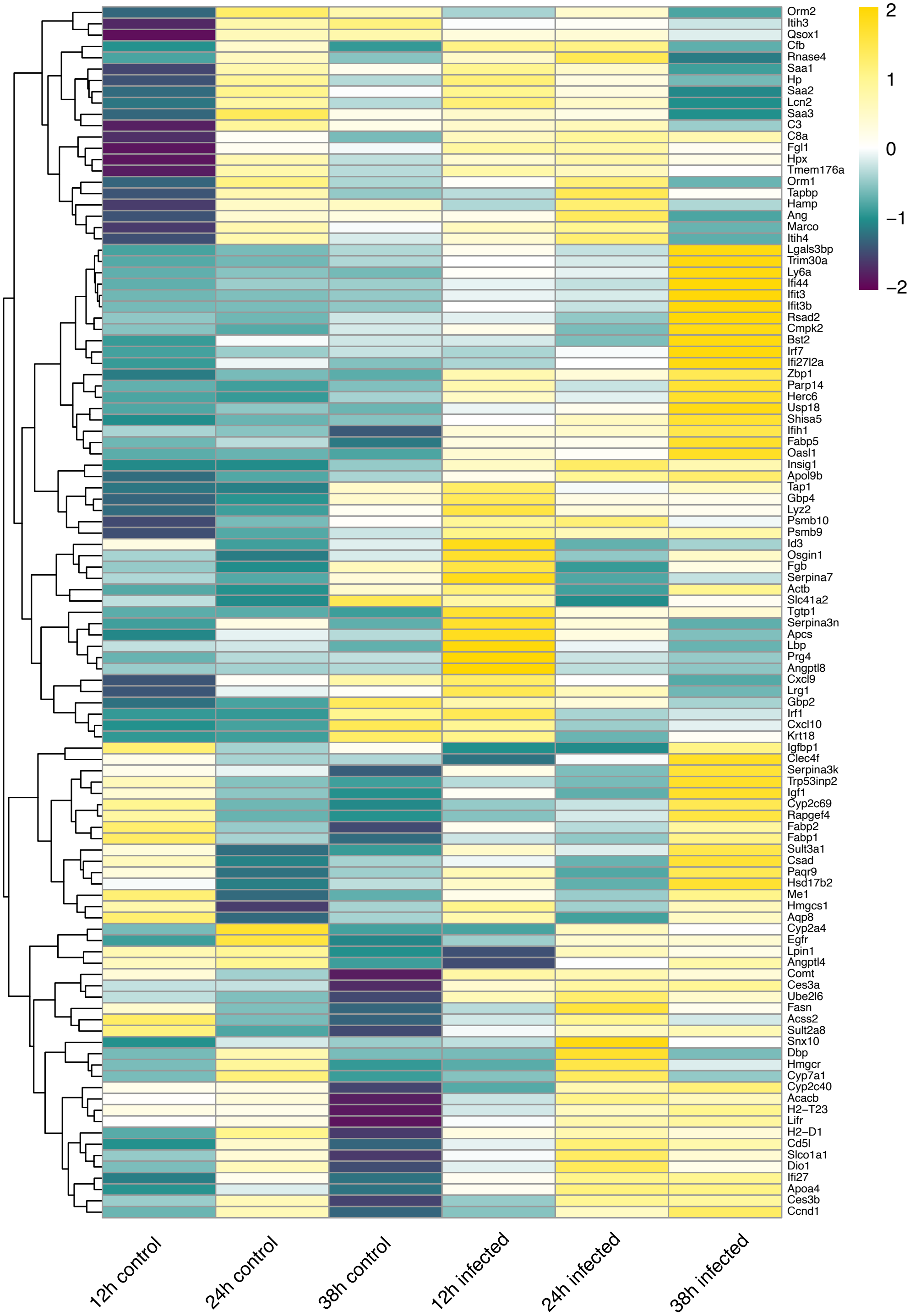


**Supplementary figure 7: Unique differentially expressed genes between SGC and infected sections across ST sections of all timepoints (12,24 and 38 hpi).** Differentially expressed genes between SGC and infected sections were determined for each individual timepoints and average expression of uniquely expressed genes across spots were visualized in a heatmap and clustered hierarchically according to gene expression similarity. Gene expression was grouped by condition and gene expression values depicted as a color scale from low expression (dark purple) to high expression (yellow).

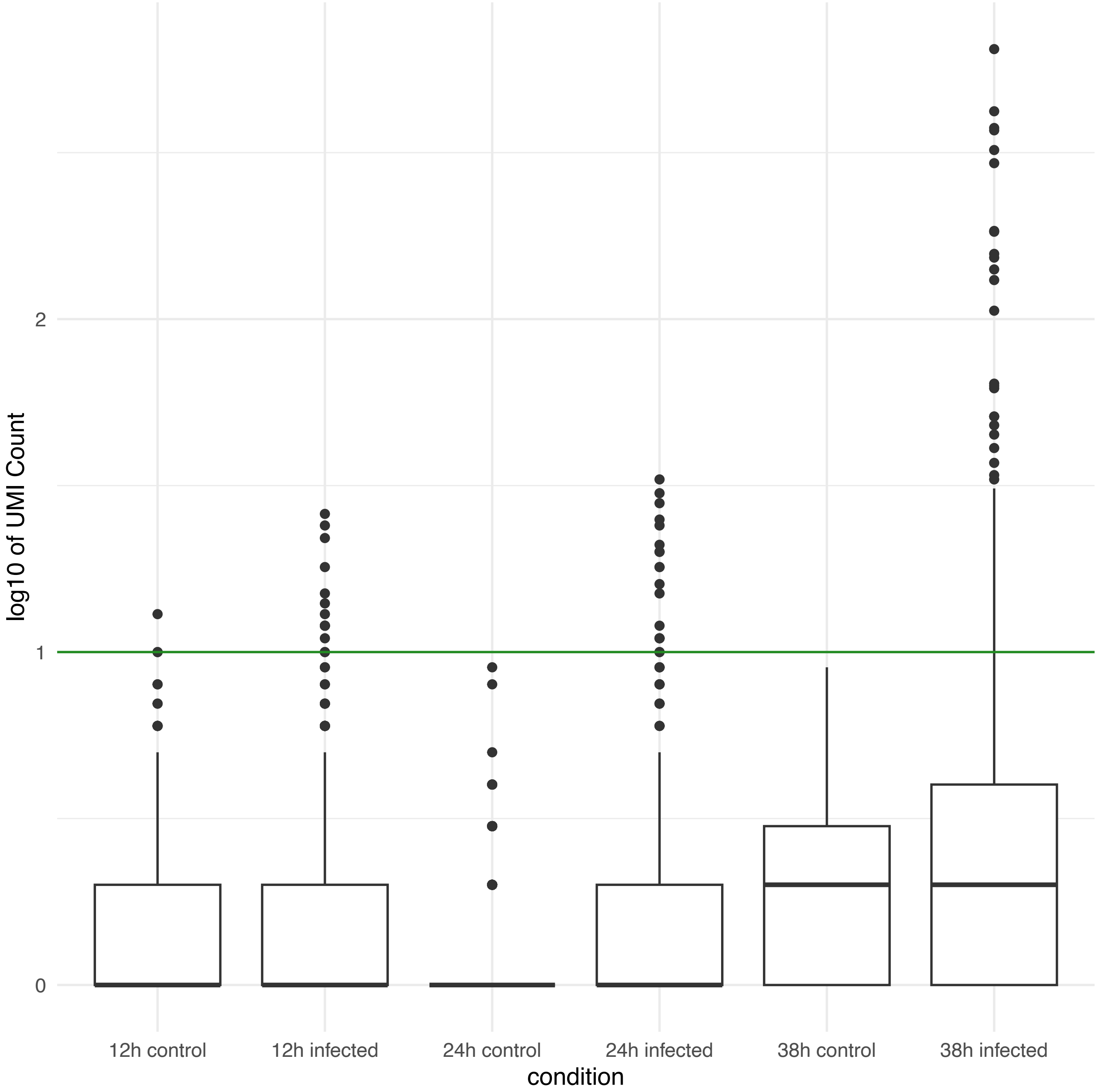


**Supplementary figure 8: Logarithmic count of unique *P. berghei* transcripts across conditions.** Boxplot of the total RNA (unique molecular identifier (UMI)) was determined across all spots and conditions. The threshold of a a high log10 count of 1 is shown in green. Upper and lower quantiles are shown as white boxes, which are divided by a black line denoting the respective median log10 count. Each point denotes a spatial position in the tissue.

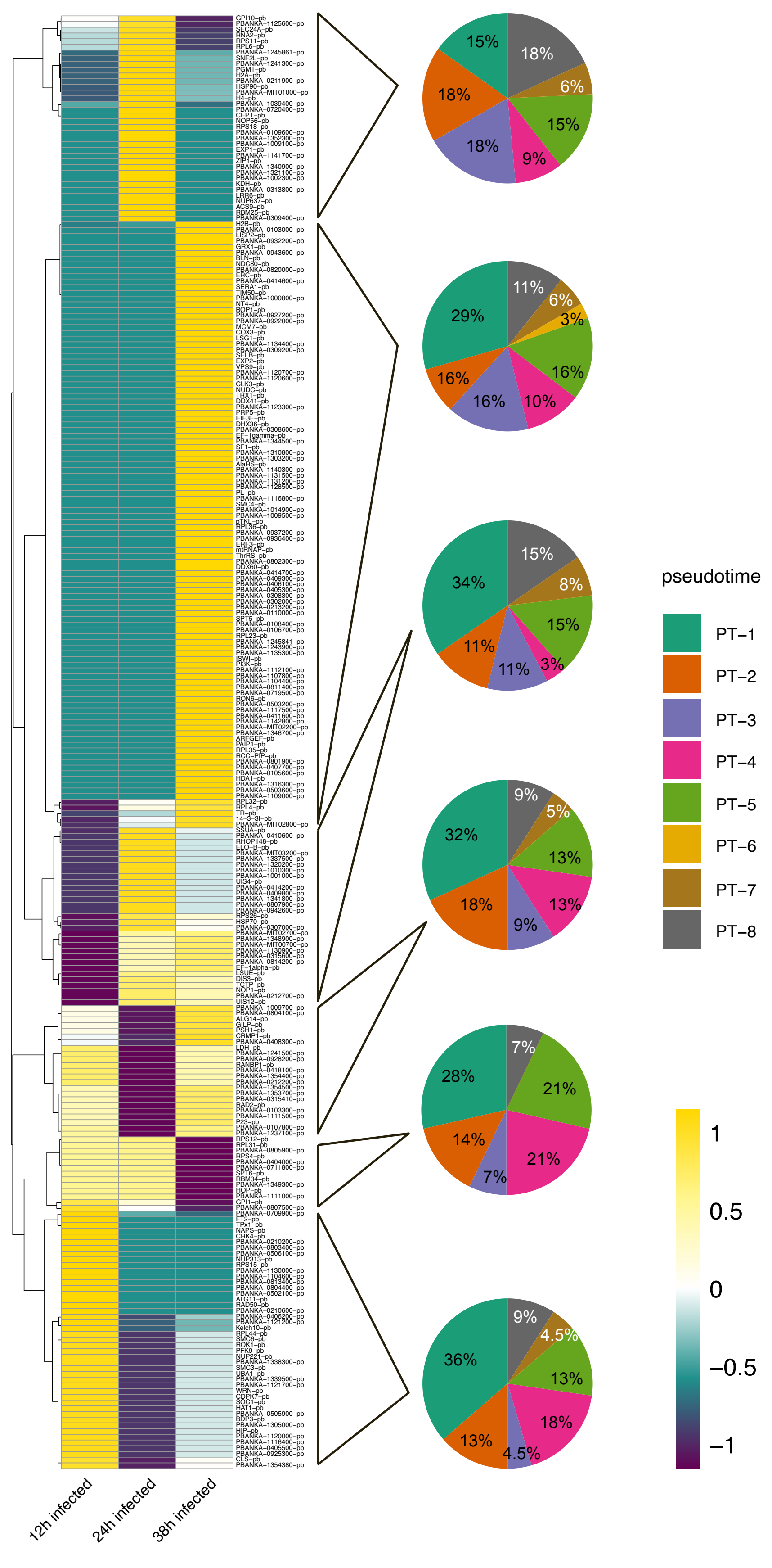


**Supplementary figure 9: Time-dependent expression of *P. berghei* transcripts.** Differentially expressed *P. berghei* genes between investigated timepoints. Average expression of genes are shown in a color gradient from low expression (dark purple) to high expression (yellow). Genes are clustered hierarchically based on similar expression patterns and grouped by infection timepoint. The 6 largest clusters of similiar expression were compared to pseudotime analyses of single hepatocyte infecting parasites by *Afriat et al., 2022.* Pie-charts show percentage of overlapping DE parasite genes with genes expressed across previously identified pseudotime bins (PT-1 to PT-8).

a

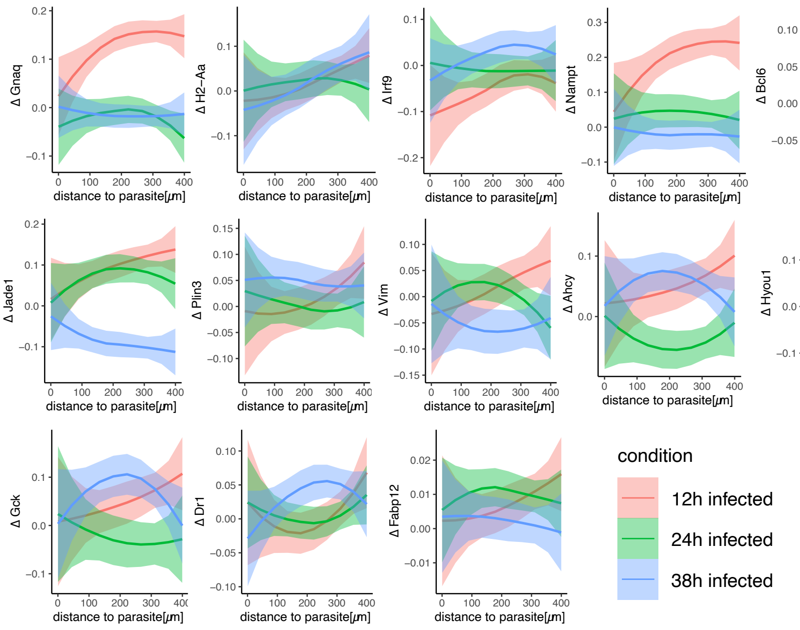


b

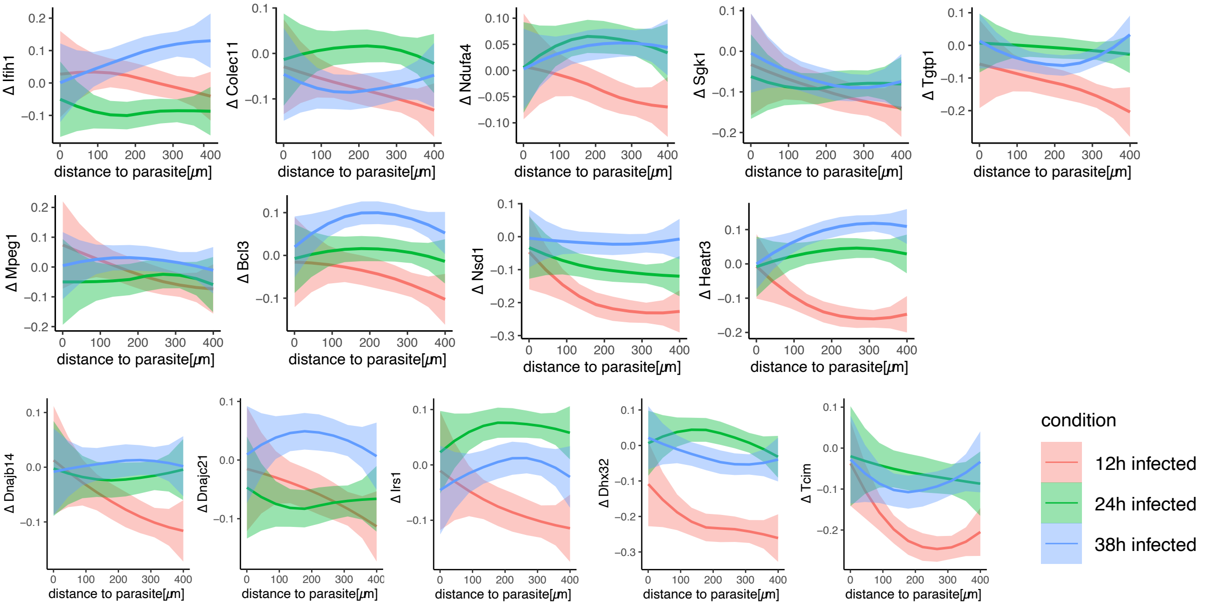


**Supplementary figure 10: Host genes exhibiting correlation between parasite distance and gene expression 12 hpi. a)** Selected host genes showing negative correlation between distance to the parasite and gene expression. Expression values are shown as average difference in gene expression (Δ) along the spatial axis. Ribbons show the standard error. Only genes expression a difference of Δ ≥ 0.1 were selected. **b)** Selected host genes with negative correlation between distance to the parasite and gene expression. Expression values are shown as average difference in gene expression (Δ) along the spatial axis. Ribbons show the standard error. Only genes expression a difference of Δ ≤ -0.1 were selected.

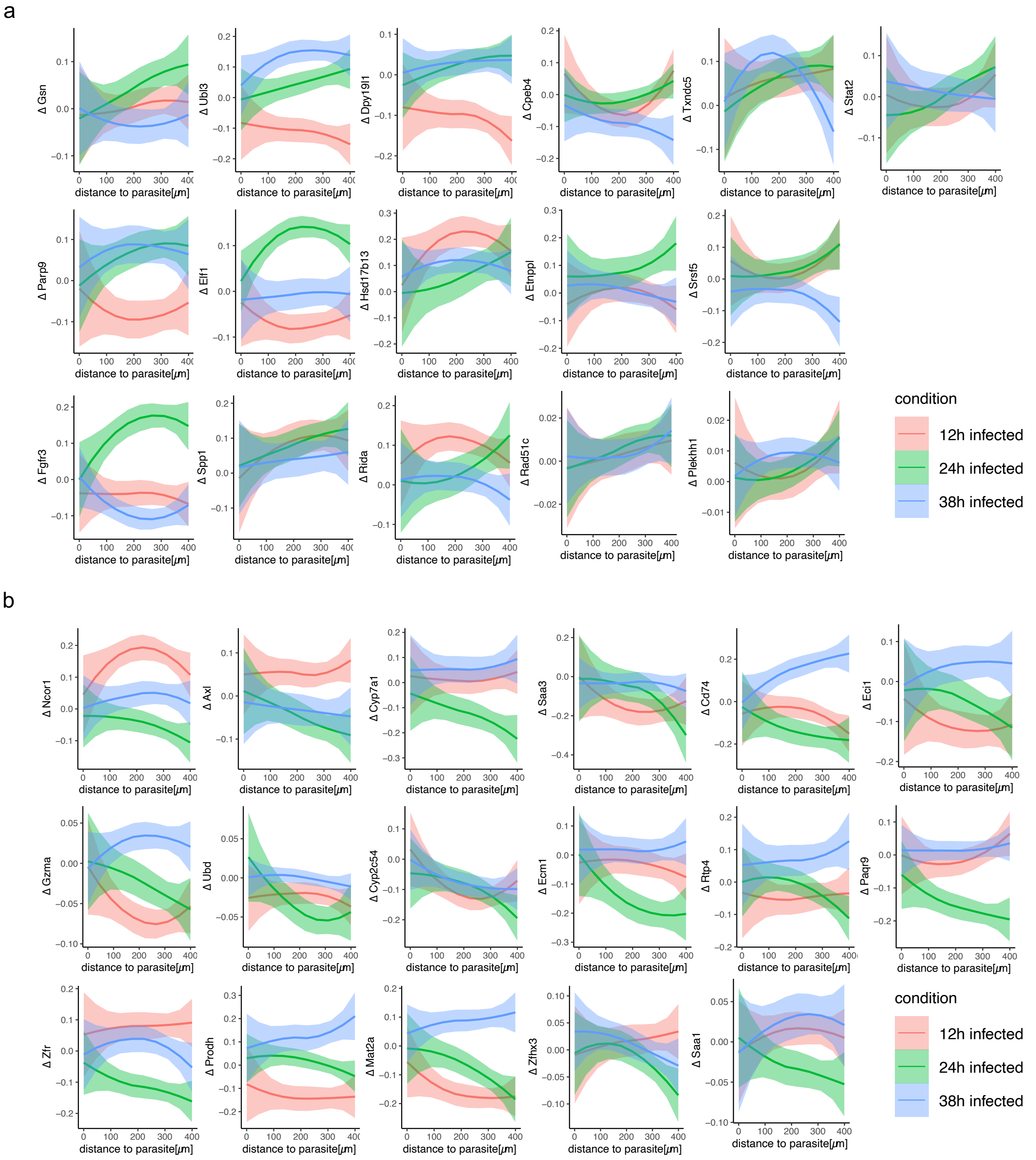


**Supplementary figure 11: Host genes exhibiting correlation between parasite distance and gene expression 24 hpi. a)** Selected host genes showing negative correlation between distance to the parasite and gene expression. Expression values are shown as average difference in gene expression (Δ) along the spatial axis. Ribbons show the standard error. Only genes expression a difference of Δ ≥ 0.1 were selected. **b)** Selected host genes with negative correlation between distance to the parasite and gene expression. Expression values are shown as average difference in gene expression (Δ) along the spatial axis. Ribbons show the standard error. Only genes expression a difference of Δ ≤ -0.1 were selected.

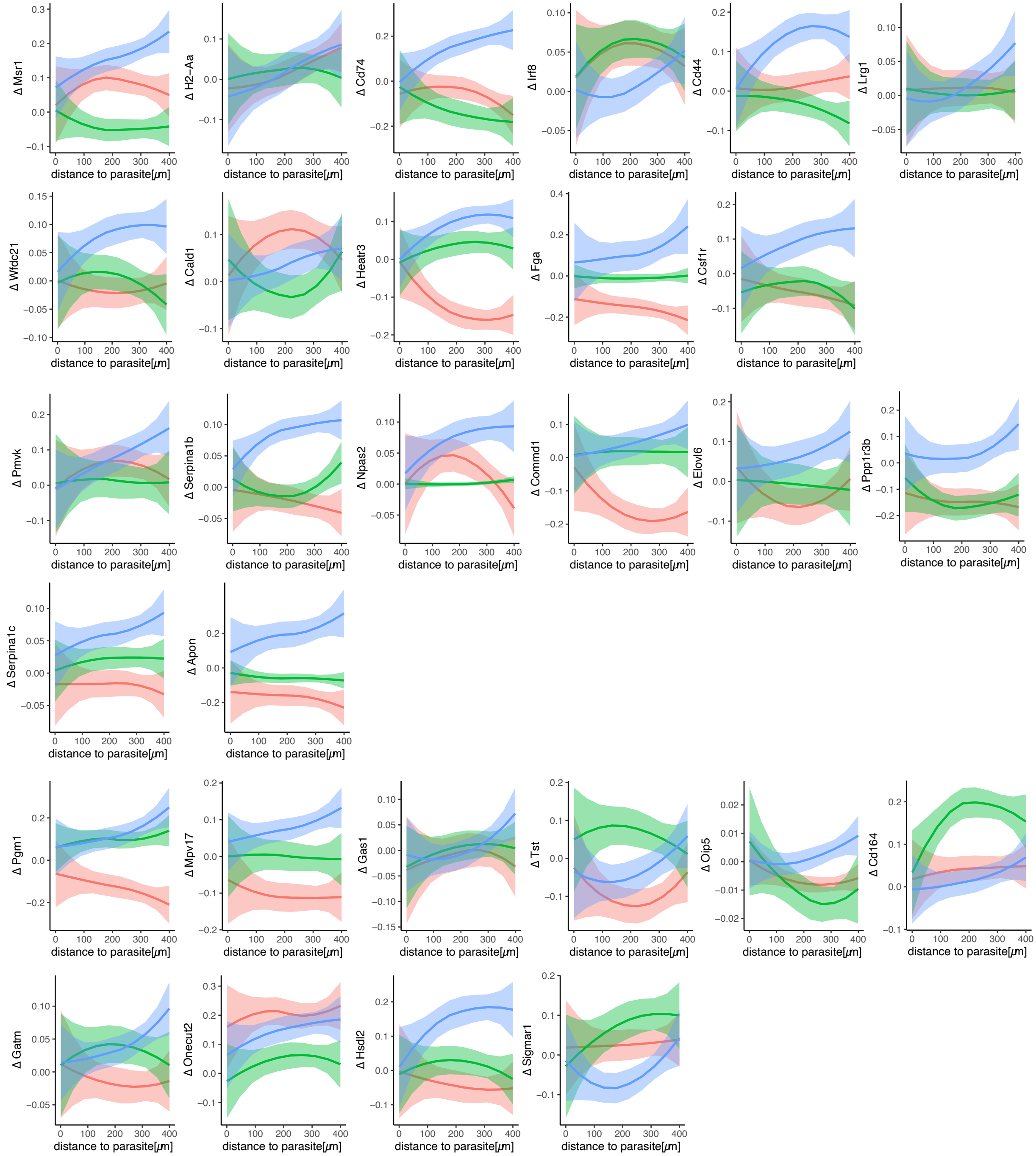


**Supplementary figure 12: Host genes exhibiting positive correlation between parasite distance and gene expression 38 hpi.** Selected host genes showing negative correlation between distance to the parasite and gene expression. Expression values are shown as average difference in gene expression (Δ) along the spatial axis. Ribbons show the standard error. Only genes expression a difference of Δ ≥ 0.1 were selected.

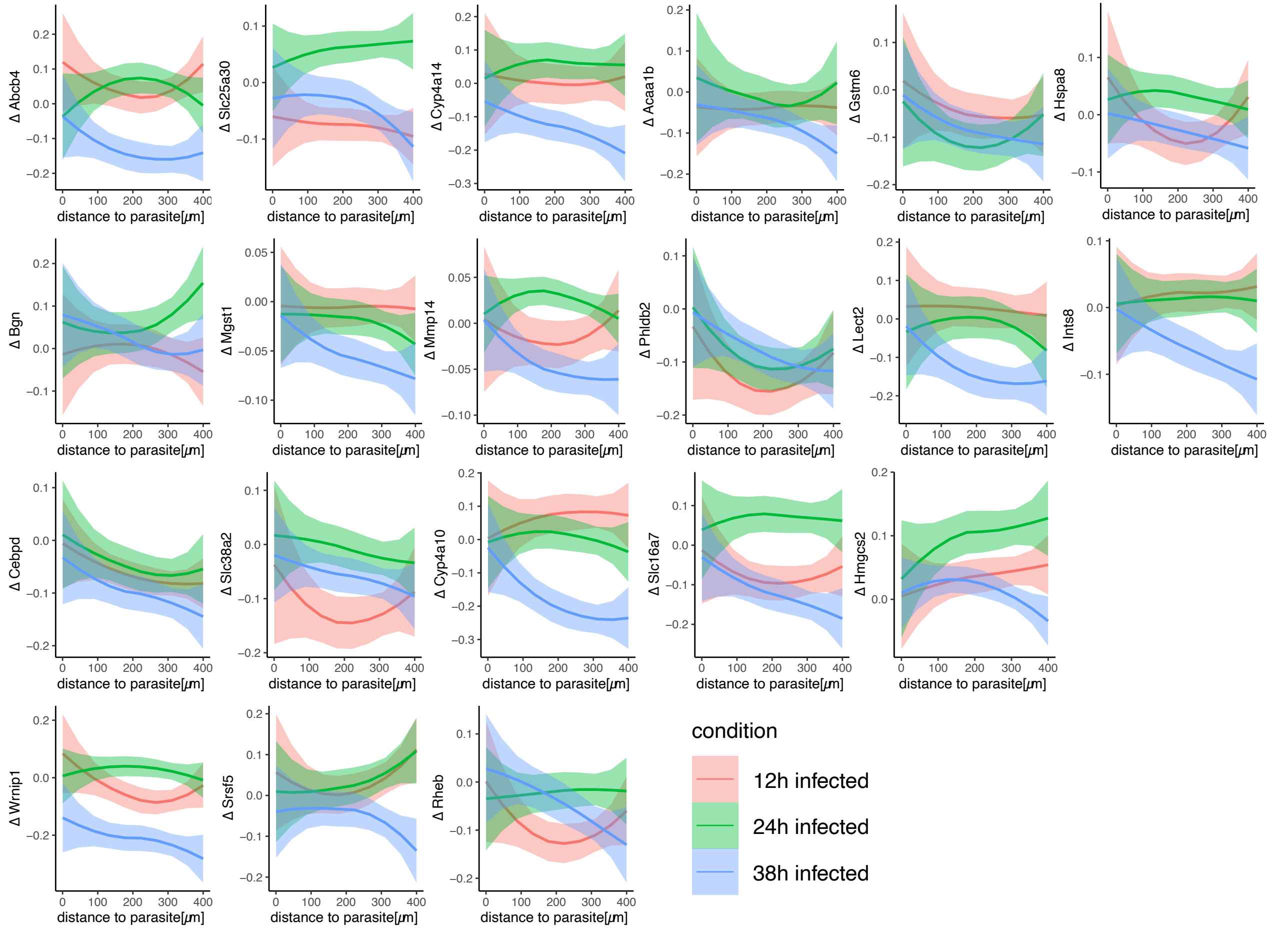


**Supplementary figure 13: Host genes exhibiting negative correlation between parasite distance and gene expression 38 hpi.** Selected host genes with negative correlation between distance to the parasite and gene expression. Expression values are shown as average difference in gene expression (Δ) along the spatial axis. Ribbons show the standard error. Only genes expression a difference of Δ ≤ -0.1 were selected.

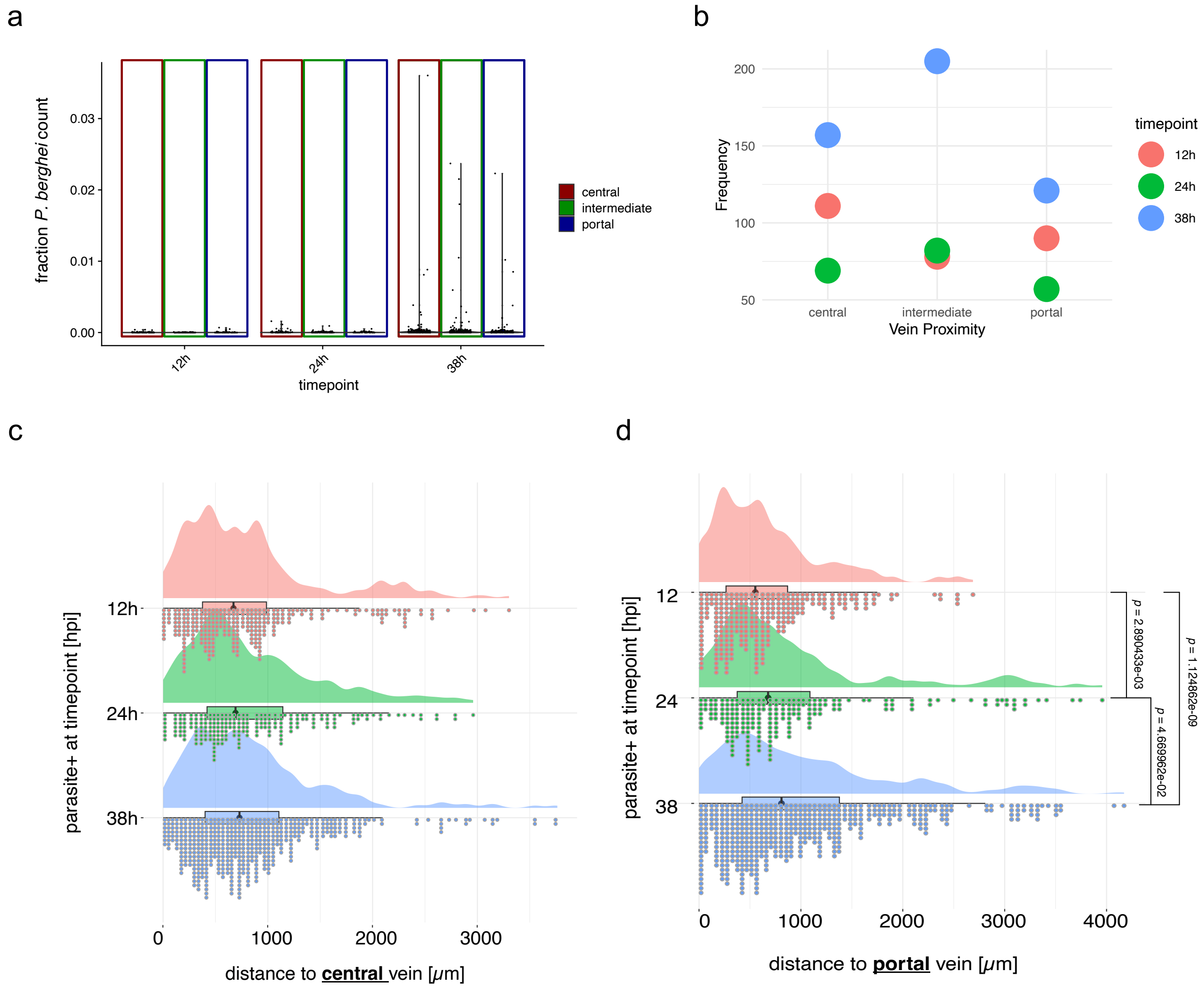


**Supplementary figure 14: Relationship between parasite and liver zonation in spatial data. a)** Fraction of *P. berghei* UMIs within 400μm of an annotated central (red) or portal (blue) neighborhood as well as fraction in intermediate (green) regions (> 400 μm of any vein). **b)** Frequency parasite annotations in close proximity to a central or portal vein as well as intermediate areas (defined as in **a)**) across infection timepoints (12 (red), 24 (green) or 38 (blue) hpi). **c)** Parasite annotations across the distance (0-3000μm) to central veins for each timepoint (12 (red), 24 (green) or 38 (blue) hpi). No significant differences were observed between timepoints **d)** Parasite annotations across the distance (0-4000μm) to portal veins for each timepoint (12 (red), 24 (green) or 38 (blue) hpi). Significant differences were obsereved between all timepoints indicated by p-values (*p*) below 0.05.

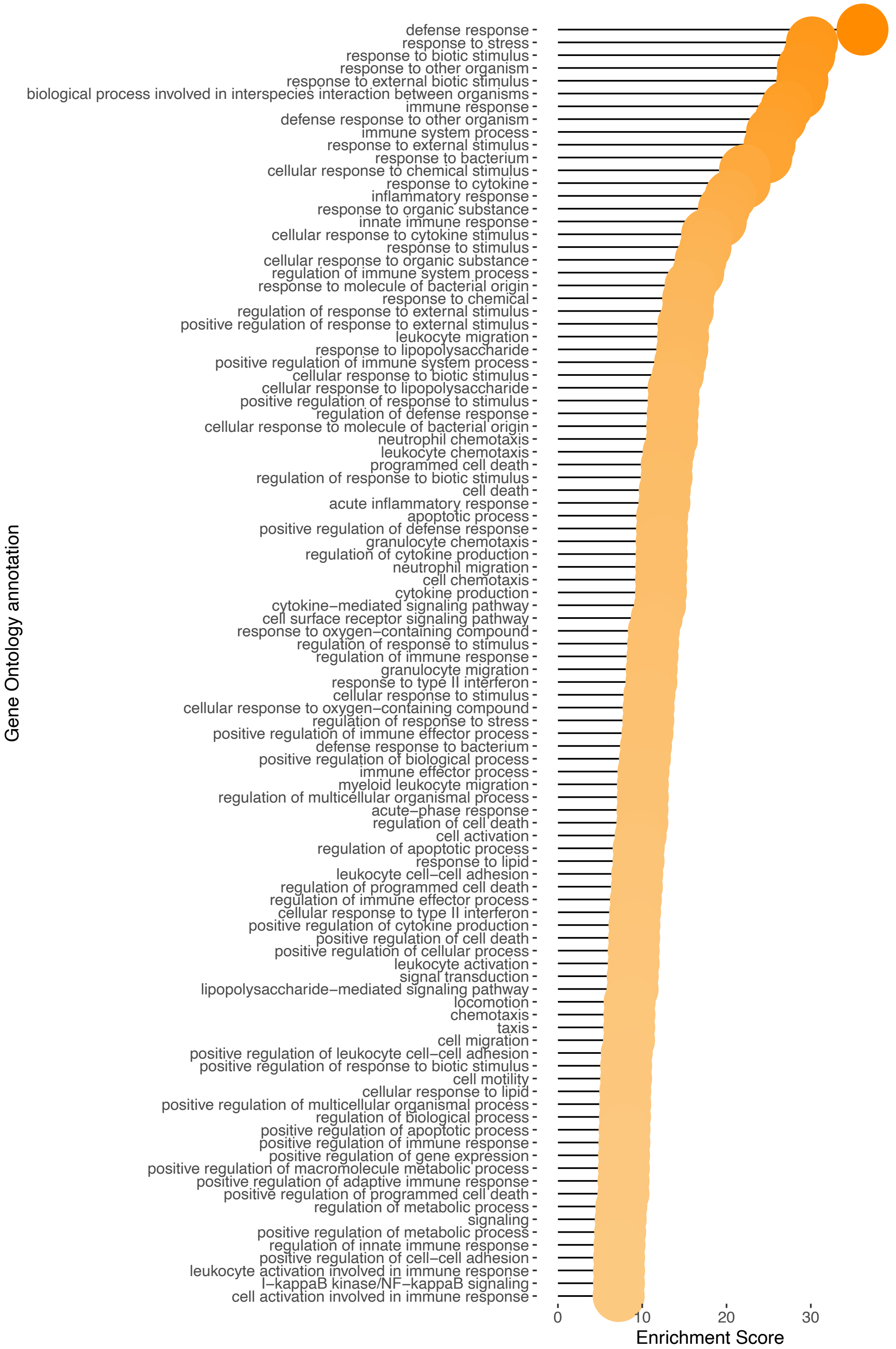


**Supplementary figure 15: Gene ontology enrichment of genes exhibiting negative correlation with IHS distance.** Gene ontology annotations showing enrichment in areas in close proximity to IHS positions. High enrichment score are displayed in darker orange.

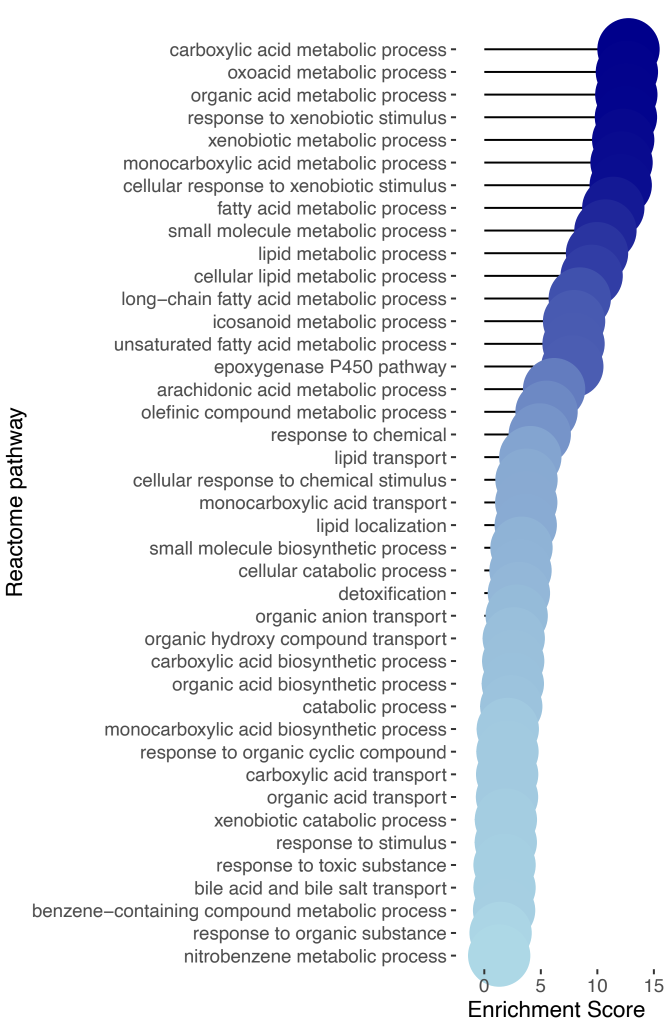


**Supplementary figure 16: Gene ontology enrichment of genes exhibiting positive correlation with IHS distance.** Gene ontology annotations showing enrichment in areas in periphery of IHS positions. High enrichment score are displayed in darker blue.

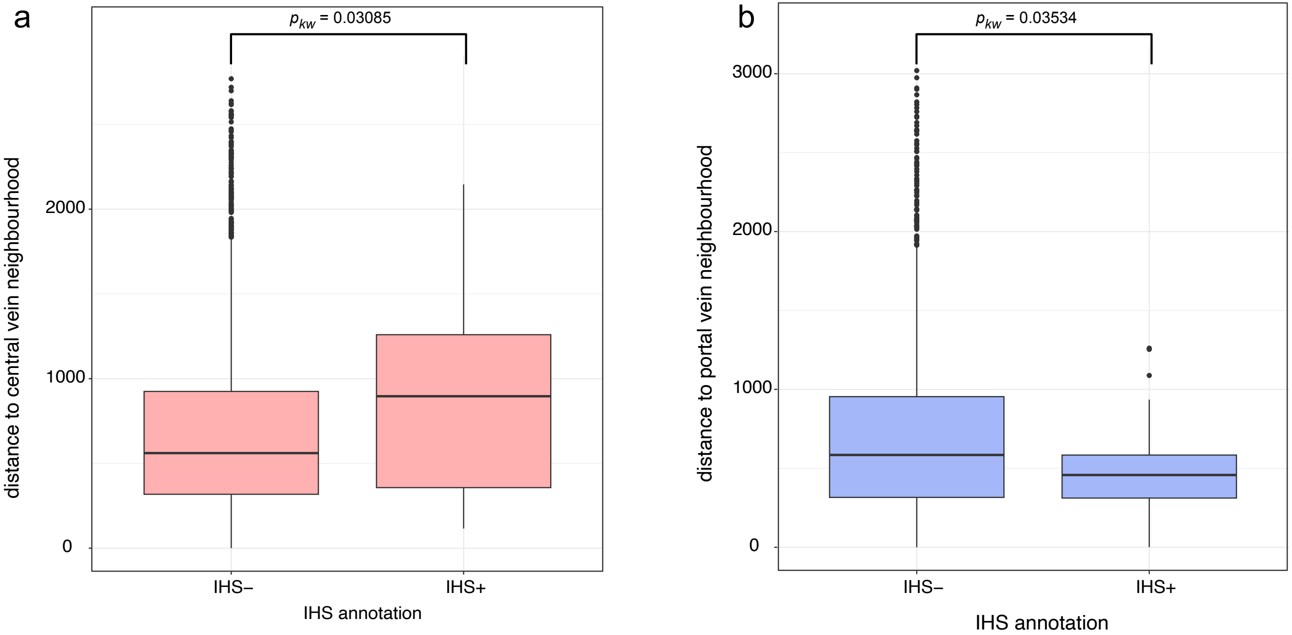


**Supplementary figure 17: Relationship between IHS position and lobular axis in spatial data. a)** average distance of spatial positions without IHS annotation (IHS-) and with IHS annotation (IHS+) to central vein neighborhood (red). Significance was tested using a non-parametric kruskal-wallis test. **b)** average distance of spatial positions without IHS annotation (IHS-) and with IHS annotation (IHS+) to portal vein neighborhood (blue). Significance was tested using a non-parametric kruskal-wallis test.

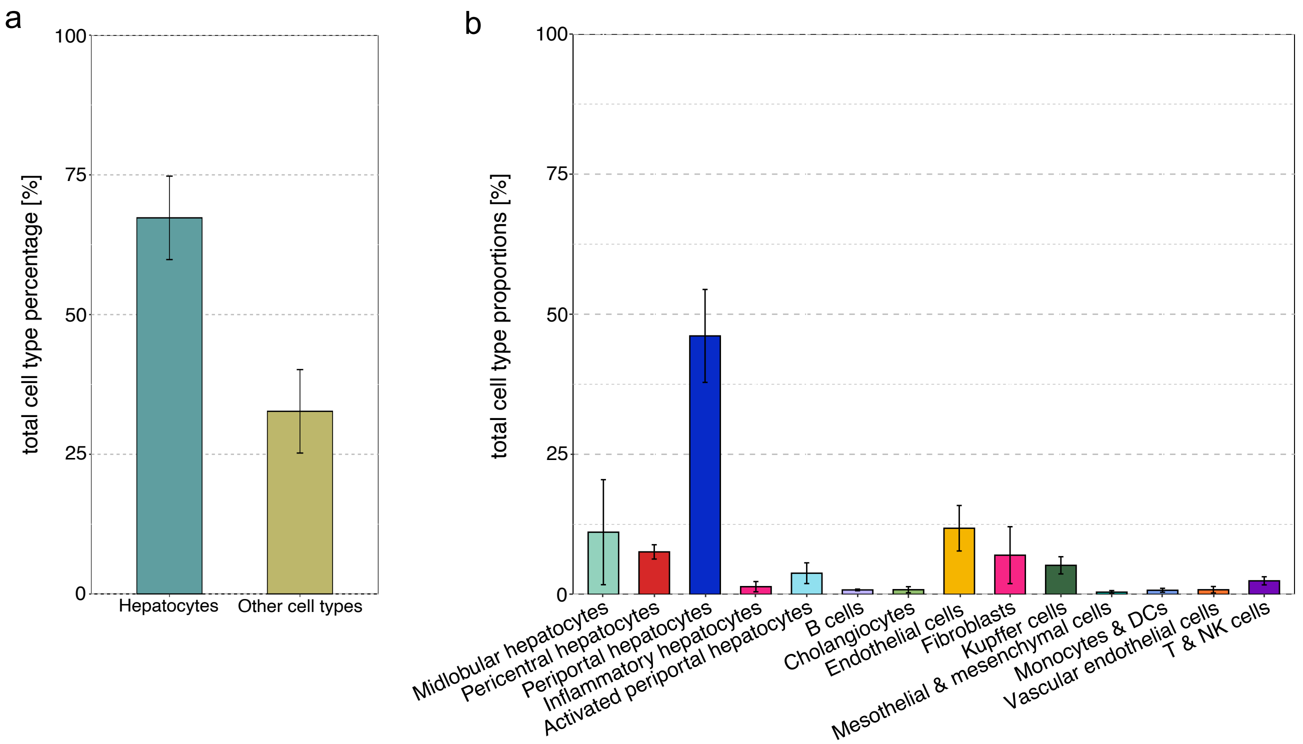


**Supplementary figure 18: Proportions of cell types across all samples. a)** average proportion of hepatocytes across all conditions and time points compared to the average proportion of all other cell types identified. **b)** average proportion of all cell types across all conditions and time points. Error bars indicate the standard error of the mean across all samples.

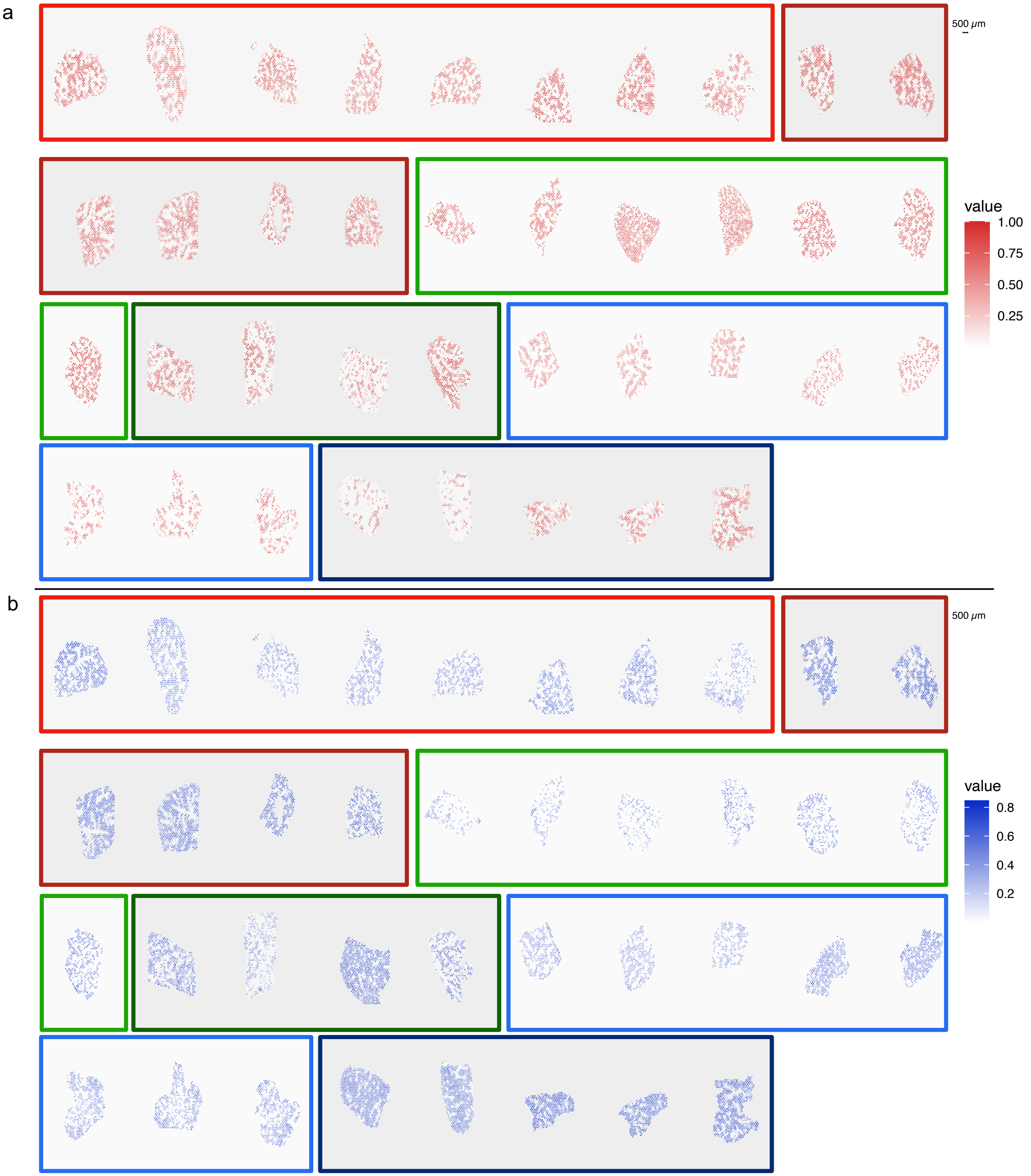


**Supplementary figure 19: Proportions of periportal and pericentral hepatocytes across all samples.** Visualization of **a)** pericentral and **b)** periportal cell type proportions across spatial positions of sections generated by ST protocol. 12h timepoints are highlighted in red, 24h in green and 38h in blue boxes. SGC sections are implicated by darker shade of the box.

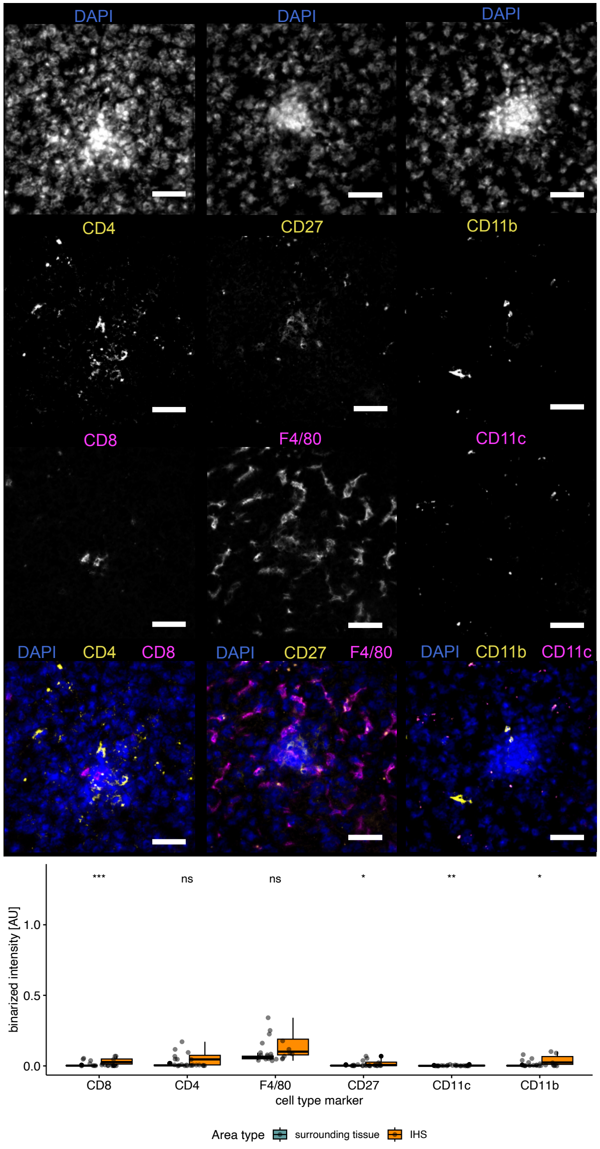


**Supplementary figure 20: Immunofluorescent staining of immune cells in infected liver tissue 12 hpi.** Representative images of staining of surface markers of two subsets of T cells (CD4, CD8), two subsets of macrophages (F4/80) and dendritic cells, monocytes and macrophages (CD11b, CD11c) as well as B cells, naive T, Treg and NK cells (CD27, CD11b) on 3 consecutive sections and inflammatory hotspots (IHSs). Images were analyzed using the same setting for each surface marker and followed by adjusting brightness and contrast for visualization (see methods for details). CD4, CD27 and CD11b are shown in yellow, CD8, F4/80 and CD11c in magenta, and DNA counterstain (DAPI) in blue in the composite (bottom). The scalebar indicates 50 µm. Control sections after 12 hpi did not show presence of IHSs.

Boxplots show binarized signal for each cell type marker indicating higher presence or absence of cells in IHS compared to the surrounding tissue for all measured IHSs. Significance levels (wilcoxon-rank-sum test, holm-corrected) are indicated by asterisks above each cell type marker.

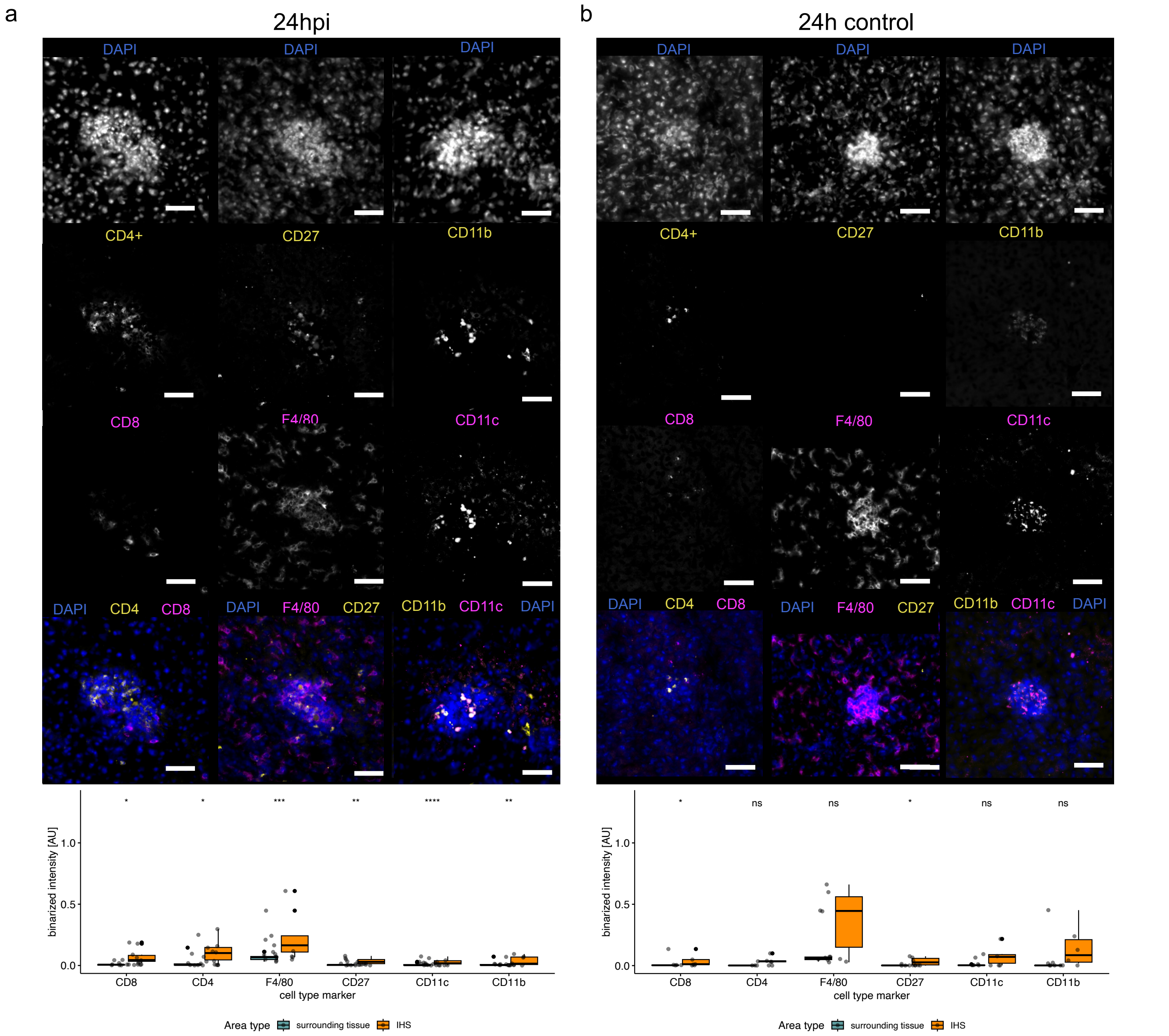


**Supplementary figure 21: Immunofluorescent staining of immune cells in infected liver tissue 24 hpi.** Representative images of staining of surface markers of two subsets of T cells (CD4, CD8), macrophages (F4/80) and dendritic cells, monocytes and macrophages (CD11b, CD11c) as well as B cells, naive T, Treg and NK cells (CD27, CD11b) of 3 consecutive sections and inflammatory hotspots (IHSs) in **a)** infected and **b)** control sections. Images were analyzed using the same setting for each surface marker and followed by adjusting brightness and contrast for visualization (see methods for details). CD4, CD27 and CD11b are shown in yellow, CD8, F4/80 and CD11c in magenta, and DNA counterstain (DAPI) in blue in the composite (bottom). The scalebar indicates 50 µm. Boxplots show binarized signal for each cell type marker indicating higher presence or absence of cells in IHS compared to the surrounding tissue for all measured IHSs. Significance levels (wilcoxon-rank-sum test, holm-corrected) are indicated by asterisks above each cell type marker.

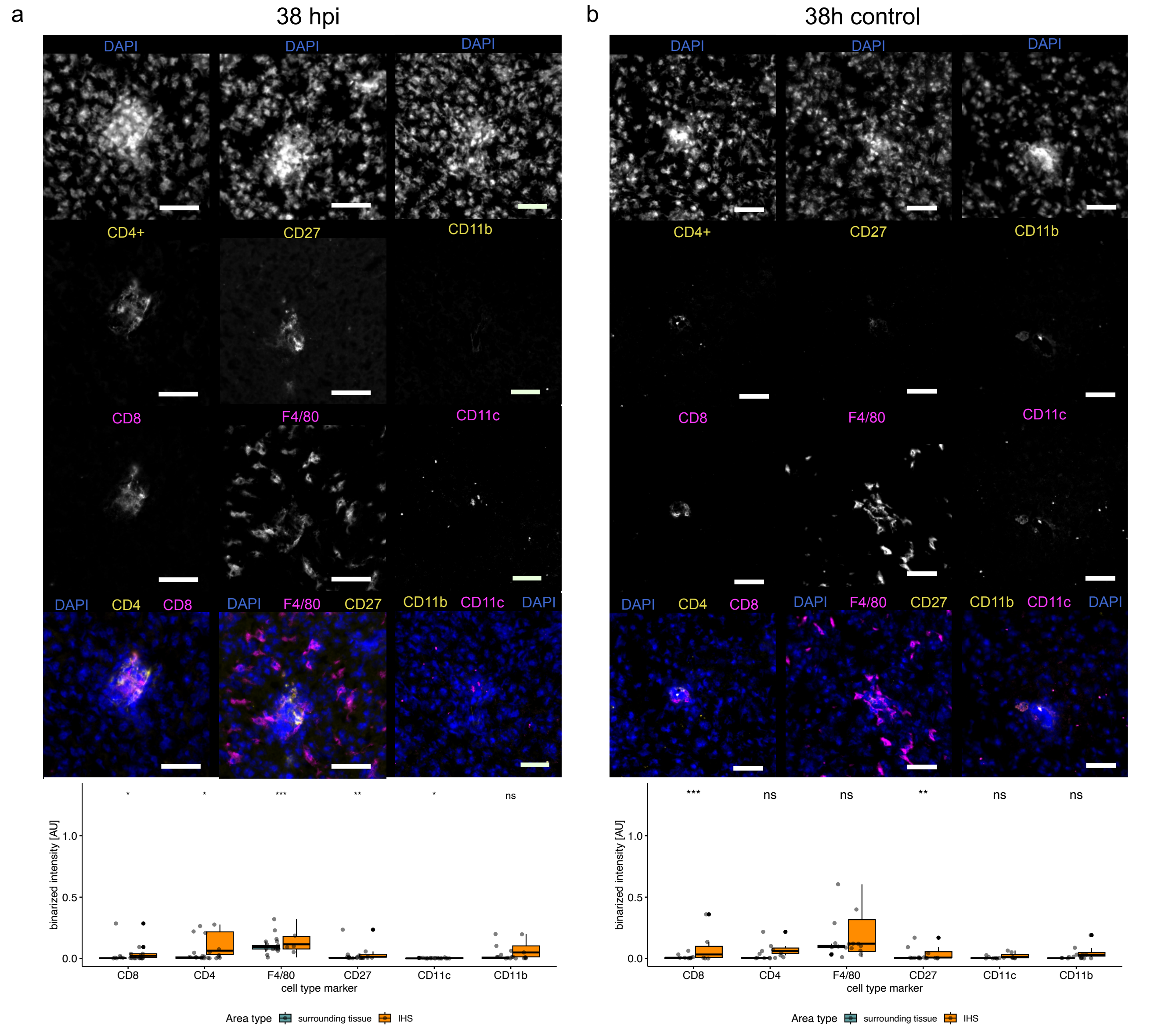


**Supplementary figure 22: Immunofluorescent staining of immune cells in infected liver tissue 38 hpi.** Representative images of staining of surface markers of two subsets of T cells (CD4, CD8), macrophages (F4/80) and dendritic cells, monocytes and macrophages (CD11b, CD11c) as well as B cells, naive T, Treg and NK cells (CD27, CD11b) of 3 consecutive sections and inflammatory hotspots (IHSs) in **a)** infected and **b)** control sections. Images were analyzed using the same setting for each surface marker and followed by adjusting brightness and contrast for visualization (see methods for details). CD4, CD27 and CD11b are shown in yellow, CD8, F4/80 and CD11c in magenta, and DNA counterstain (DAPI) in blue in the composite (bottom). The scalebar indicates 50 µm. Boxplots show binarized signal for each cell type marker indicating higher presence or absence of cells in IHS compared to the surrounding tissue for all measured IHSs. significance levels (wilcoxon-rank-sum test, holm-corrected) are indicated by asterisks above each cell type marker.
